# The effects of Hurricane Otto on the soil ecosystems of three forest types in the Northern Zone of Costa Rica

**DOI:** 10.1101/2020.03.19.998799

**Authors:** William D. Eaton, Katie M. McGee, Kiley Alderfer, Angie Ramirez Jimenez, Mehrdad Hajibabaei

## Abstract

Hurricanes rapidly deposit large amounts of canopy material onto tropical forest floors, stimulating metabolic processes involved in the decomposition of these materials and production of N and C resources into the food web. However, little is known about the effects that hurricanes have on specific soil microbial taxa or functional groups involved in these processes. The objectives of this study were to determine how Hurricane Otto influenced three different tropical forest soil ecosystems within the first 8 months after causing damage to a tropical forest by assessing the soil C and N factors and how the soil bacterial and fungal community compositions differed before and after the hurricane. Soil samples were collected from five 2000 m^2^ permanent plots in Lowland, Upland and Riparian forest systems within the same area in the Northern Zone of Costa Rica. Standard methods were used to measure the amounts Total N, NO_3_^-^, NH_4_^+^, Total organic C, and Biomass C, while Illumina MiSeq methods were used to generate bacterial and fungal sequences. All data were analyzed using univariate and multivariate statistical methods. Using this “before and after” study design, it was found that the levels of the inorganic N and Biomass C were greater in the Post-Hurricane soil samples. The mean proportion of DNA sequences of complex C degrading/lignin degrading, NH_4_^+^-producing, and ammonium oxidizing bacteria, and the complex C degrading/wood rot/lignin degrading and ectomycorrhizal fungi also were greater in the Post-Hurricane soils. We also provide evidence that the excessive amounts of canopy leaf litter and woody debris deposited on the forest floor during Hurricane Otto appears to be selecting for genera that become more dominant Post-Hurricane, perhaps because they may be better able to rapidly process the newly deposited C and N-rich canopy material. This is a rare “before and after” a natural hurricane design that warrants further monitoring.

## Introduction

The occurrence of tropical hurricanes are thought likely to be increasing in frequency in the future [1], [2]. These disturbances alter the forest vegetation composition and productivity, through tree damage [3]. Hurricanes cause rapid deposition of significantly large amounts of leaf litter and woody debris from the canopy to the forest floor [4], [5], [6], [7], which then results in short to long-term changes to the forest structure and forest ecological processes [4]. These processes are critical to forest recovery, and are associated with both the rapid increase in the carbon (C) and nitrogen (N) nutrients [8], [9], [10], [11] and the subsequent influences to the soil microbiota [12], [13].

There have been studies on the effects of hurricane-induced litter fall and enhanced soil nutrients on soil processes (especially decomposition) that have shown the patterns observed are a function of the characteristics of the specific hurricane, and that of the forest before the hurricane hit, making it difficult to predict specific effects from the hurricane-induced dropping of canopy material [8]. Moreover, much of the leading work has been conducted under hurricane simulation experiments in which canopy components are cut and dropped to the forest floor (for example: [6] [9] [14] [15] [16]), which leads to the important question of “what happens under real conditions and are they different?”. Clearly, there is much work to be conducted, but doing work in the same forest location before and after a hurricane is truly a serendipitous opportunity. This paper describes just such a serendipitous opportunity to study the effects of a hurricane on the same tropical forest soils before and after a hurricane in three different habitats within a Costa Rican Tropical forest.

Lugo [8] suggests that there are shorter-term Immediate Effects that a hurricane has on a tropical forest of 0 to 3 years, and possible longer-term Intermediate Responses of 0 to 20 years. In field experiments in which canopy material was cut and dropped to the forest floor to simulate the effects of a hurricane, nutrient pulses of more labile forms of C, N and phosphorus derived from the younger leaf litter from the canopy were processed in the first year post deposition of material [4][6] [9] [10] [16] [17]. Following this, Liu et al ([10]) showed the initial pulse of labile organic C and nutrient turnover rate in the forest soil lasted from about the first week to about 2 years. They also showed there was an elevation of the soil microbial biomass that lasted from week 1 until at least 120 weeks after deposition of the canopy material. The hurricane-induced deposition of course woody debris (CWD) from the canopy caused an increased immobilization of the nutrients, resulting in a decreased availability of these for the food web [11] [18] [19] [17] [20], a slowing down decomposition activities in the forest soils [9].

Soil bacteria are involved in C and N processes associated with early stages of soil succession or recovery from disturbance, such as decomposition of less complex organic C, N-fixation, and ammonium oxidation [21] [22]. Major forest disturbances, such as hurricanes, that alter the quantity and quality of plant nutrient input into the forest soil, the concentrations of critical soil substrates, and the pH and temperature levels in the soil will influence the composition of the soil bacterial communities and the rates and outputs from these different C and N processes [23] [24] [25]. However, due to their critical role in the C and N processes, these same bacterial groups are expected to be some of the first biota to re-establish in the soils during the early recovery stages. The soil fungi are important in later soil successional stages and later stages during the soil recovery processes. This is due to their abilities to decompose complex organic substrates that develop during earlier stages of soil succession or recovery more efficiently than bacteria, leaving behind more recalcitrant residues, and enhancing the soil organic C matter [26] [27] [28]. These fungi require the N and C inputs from the earlier bacterial activities in order to produce the enzymes necessary for activities such as wood rot, degradation of lignin, and decomposition of other complex forms of organic C [26] [27] [28].

Some experimental work has been conducted simulating the hurricane-induced deposition of canopy leaf litter and CWD in a tropical forest, in order to identify the potential influences that this fallen canopy material might have on the forest floor C, N, and to a lesser extent, the soil microbial communities [14][15]. These studies have shown that there are differences in overall microbial community structure between forest floor soils with leaf litter and CWD deposited on them as compared to control sites. These are extremely important studies that have helped to develop some basic information on the potential activity of different soil processes that might occur post-hurricane, however, they do not identify how any specific microbial taxa or specific microbial functional groups within the soils might be affected by the addition of the vegetation material Cantrell et al. [15] found differences in the overall bacterial and fungal communities, bacterial richness, and bacterial and fungal diversity between the soils with leaf litter and CWD deposited on them and untreated forest floor soils, using ester link fatty acids methyl ester (EL-FAME) and terminal restriction fragment length polymorphism (TRFLP) analyses. Lodge et al [14] found differences in the amount of fungal connections present in leaf litter between these treated and untreated soils using visual observations. Others have shown an increases in soil C, microbial biomass C and N between sites with experimentally deposited leaf litter and CWD, also between these sites by season, and beneath and in close proximity to the larger CWD from the experimental vegetation deposition as compared to the soil more distant to the CWD [9] [15]. Another study showed there was an initial (1 week post experimental deposition of leaf litter and CWD) to longer-term (120 weeks post-deposition) increase in microbial biomass C occurring in the leaf litter of the forest floor during the experiment [10]. In a review by Lugo [8], the idea that the extensive defoliation of young leaf material and its deposition on the forest floor also is thought to enhance early N cycle activity, and perhaps results in changes in the associated soil microbial communities. In fact, McDowell et al [29] found that riparian soils had increased levels of nitrate and ammonium following hurricane-induced deposition of leaf litter, which was absorbed from the soils by the vegetation within the first few years. However, here too no information was provided on changes in the microbial communities associated with these changes in inorganic N.

What appears to be missing from these studies, and the literature in general, are the classic “before and after” studies in which forest soils are analyzed for differences in C and N nutrients, microbial biomass C, fungal and bacterial community composition of the overall genera, and in the composition of different important microbial functional group taxa between the same forest plots before and after a naturally occurring hurricane, instead of a simulation. On November 24, 2016, Hurricane Otto hit the forests in the Northern Zone of Costa Rica providing the serendipitous opportunity to conduct just such a before and after study in the exact same long-term forest plots. These plots had been permanently established previously in three primary forest types within the Maquenque National Wildlife Refuge in the Northern zone of Costa Rica, an Upland Forest Type, Lowland Forest Type, and Riparian Forest Type, and used for various studies [29] [30] [31] [31] [32] [33] [34], as well as for the soil samples collected for a project in the summer of 2016. Upon arrival at the sites in summer 2017, we found that all plots in all three forest types had been hit hard by Otto, significantly decreasing the percent canopy cover by about 40 to 70%, and littering the forest floor with young leaf litter and large amounts of CWD. This provided us with the opportunity for the classic before and after study we report on in this article. Thus, the combination of our previous work in established permanent forest plots, and Hurricane Otto provide a very rare opportunity to study the soil microbial communities and C and N cycle metrics within 3 primary forest habitats before and only 8 months after Hurricane Otto, within the exact same long-term plots. This is still within the timeframe for evidence of early effects of a hurricane [8].

The goal of this study was to determine how a hurricane influences the soil ecosystem conditions within the first 8 months after causing damage to a tropical forest. To do this, our aim was to determine how soil C and N factors and associated bacterial and fungal community compositions differed before and after a hurricane caused serious damage to three different types of tropical forests within a primary forest region (Upland, Lowland, and Riparian) in the Northern Zone of Costa Rica. For this work, five broad questions were asked: (1) How do the soil C and N-cycle metrics, C biomass development, and the efficient use of the soil organic C as the Microbial Quotients [35], [36] differ between primary forest soils before and after the hurricane? (2) Are there differences in the overall bacterial and fungal community compositions between these soils? (3) Are there differences in the composition of several critical functional groups important to the N and C-cycle activity and development of the Microbial Quotient between the soils? (4) Which soil environmental variables (of the C and N cycle, biomass and Microbial Quotients metrics) appear to shape, or drive the soil microbial community structures? Based on the previous literature [4-11], we predicted that the likely deposition of fresh leaf litter and excessive woody debris deposited on the forest floor during Hurricane Otto would result in increased levels of inorganic N and C biomass, as indicators of enhanced microbial activity involved in the decomposition and incorporation of the extra C and N material into the forest biota, and would be associated with an increase in the abundance of the microbes associated with these activities.

## Methods

### Study Site

In 2001, the Costa Rican Ministry of the Environment and Energy creating the San Juan-La Selva Biological Corridor (SJLBC) to connect six protected areas into a single integrated biological unit of 1,204,812 ha. for habitat connectivity and protection of biodiversity in the Northern Zone ecosystems [37]. The Maquenque National Wildlife Refuge (MNWR; 10°27’05.7’’N, 84°16’24.32’’W) was established in 2005 by the Costa Rican government to protect over 50,000 ha. of humid Atlantic lowland primary and secondary forests and other diverse ecosystems within the SJLBC. The MNWR is the core nucleus of the SJLBC as it conserves the highest percentage of forest cover and contains the most valuable habitats for biodiversity [38]. The mean annual temperature of the MNWR is 27°C, mean annual rainfall is 4300 mm, and the dominant soil type is oxisols [38]. The habitats within the MNWR used for this project were: (a) an Uplands Primary Forest, b) a Lowlands Primary Forest, and c) a Riparian Primary Forest. All 3 forests are connected as part of a large region of Primary forest within the MNWR.

### Sample Design and Soil Collection

A bulk soil approach was used for this study to compare soil characteristics at the forest habitat level, eventually using a “before the hurricane” and “after the hurricane” design. In 2010, 5 individual replicate large 50 m x 40 m plots were permanently established in these forest types for future use. The replicate plots were between 100 m and 2 km apart, and established using the standard stratified, block, systematic plot study design used in forestry studies, studies of damaged lands, and restoration ecology studies, as recommended by the American Society of Foresters (ASF; see www.forestandrange.org) and the US Environmental Protection Agency (see EPA 2002 document Guidance on choosing a sampling design for environmental data collection--2002, EPA/240/R-02/005). Soil samples were collected in August of 2016, before Hurricane Otto hit the forests of Costa Rica, and again in June of 2017, 7 months after the hurricane hit the area. It is critical to understand the differences in spatial scale when working with soil microbes as compared to “above-ground” organisms in ecology. A distance of several meters to several kilometers is considered a landscape-level separation [39] [40]. Given this and the separation of our soil sampling sites by 100 m to 2 km, the soil microbial communities within the samples collected in this study were separated by landscape-level scales, providing 5 different representative sample plots for each forest type that were true replicates and not pseudo-replicates. To ensure no cross-contamination occurred between plots, thus maintaining the integrity of true replicates, a 7.5 cm x 15 cm x 1.25 cm soil profiler was used to aseptically collect nine soil profile cores of 0-15 cm depth in each plot, using a pre-determined sampling strategy in each plot, with 70% ethanol decontamination of all gloves and tools occurring between plots. Prior to soil collection, the litter layer was carefully removed from the forest floor, exposing the upper organic layer. The nine cores collected from each plot were combined into sterile bags, providing six composite and independent replicate samples per habitat type. After collection, using sterile technique, all soil samples were passed through a sterilized 4-mm sieve at field moist conditions prior to all downstream analyses.

### Carbon and nitrogen properties

Subsamples (200g) of field moist soil from each sieved soil sample were delivered to the Center for Tropical Agriculture Research and Education (CATIÉ) Laboratories in Turrialba, Costa Rica for determination of all C and N values. The total organic C (ToC) levels were determined via dry combustion analysis of Anderson & Ingram (1993), and the Biomass C determined by standard chloroform fumigation methods. The total N mass (TN) levels were determined by the Kjeldahl method, and NH_4_^+^ and NO_3-_^-^ levels were measured from 2M KCl extracts using the spectrophotometric methods of Alef and Nannapieri [41].

### DNA extraction, sequencing, and bioinformatics

Environmental microbial DNA (eDNA) was extracted from three 0.33g replicate sub-samples for a total of 1g for each soil sample using the MoBio PowerSoil DNA Isolation Kit (MO BIO Laboratories Inc., Carlsbad, CA, USA). The concentration and purity (A_260_/A_280_ ratio) of extracted soil eDNA were determined prior to downstream analyses using a NanoDrop 1000 spectrophotometer (ThermoFisher Scientific, Waltham, MA). All of the following methods are described in detail by McGee et al. [32] [42]. Briefly, the different eDNA extracts from the soil samples were used for a 2-step PCR amplification of eDNA targeting the nuclear internal transcribed spacer (ITS) ribosomal RNA gene region for fungi [43], and also targeting the v3 and v4 of 16S ribosomal RNA gene region for bacteria and archaea [44]. All generated soil amplicons were sequenced in several Illumina MiSeq runs using a V3 MiSeq sequencing kit (FC-131-1002 and MS-102-3003). The subsequent 16S and ITS Illumina-generated sequences were processed using semi-automated pipelines, producing operational taxonomic units (OTUs), which were processed and taxonomically assigned from Phylum to Genus using the Ribosomal Database Project (RDP) classifier v2.12 [45] for bacteria and using the RDP classifier with the UNITE fungal ITS training set for fungi [46].

The number of times a specific DNA sequence (i.e., OTU or genus) appeared in a sample was converted to mean proportion of the sequences per sample [47], hereafter called the MPS. The MPS was determined for each genus within each soil subsample from the different forest types as the number of sequences of an individual genus within a soil subsample, divided by the total number of sequences within that subsample. All generated sequencing data were submitted to GenBank, accession number XXXXX. Subsets of sequences taxonomically identified to the genus level were generated by categorizing certain genera into the different functional groups of complex C degrading and lignin degrading bacteria (CCD/Lignin), bacteria producing N ammonium (NH_4_^+^) either from N-fixation or nitrite/nitrate reduction (NH_4_^+^ Producers), complex C degrading/wood rot/lignin degrading fungi (CCD/WRT/Lignin), and ectomycorrhyizal fungi (ECM). The MPS values of these groups were also determined.

### Data analysis

All data were examined by comparing the components in the soils of the 3 different forest types before and after Hurricane Otto hit Costa Rica. To address question 1, the mean values of all C and N data were examined by one-way analysis of variance (ANOVA) followed by Tukey’s HSD or Dunnett’s T3 post-*hoc* tests, as appropriate, in SPSS (v.25, Armonk, NY, USA) to determine if there were differences in these metrics across and among the different tree soil comparisons. Prior to ANOVA, the Levene’s test was performed in SPSS to determine homogeneity of the variances of the data, and the Shapiro-Wilk test was performed in SPSS to determine normality of all the data. All data had *p* values > 0.05, suggesting the use of ANOVA was appropriate.

To address question 2, all MPS values were transformed using a 4th root transformation to account for dominant and rare taxa, as recommended by Anderson et al. [48]. The 4th root transformed genera data were then converted into Bray-Curtis dissimilarity matrices in PRIMER-E v6 for multivariate analyses, which were then used to calculate the Margalef’s richness (d=(S-1)/Log(N)) using PRIMER-E v6 [48] for the total bacterial and fungal genera and for the genera associated with the three different functional groups of interest. The magnitude of the differences in the MPS values and richness levels before and after the hurricane was assessed by PRIMER-E v6 and its PERMANOVA+ add-on [48] and the Hedge’s *g* effect size calculations. Hedge’s *g* for Effect Size is recommended over Cohen’s *d* when sample sizes are < 50, as was the case in our study. An effect size of 0.5 represents a medium magnitude effect, and ≥ 0.8 a large magnitude effect. Effect sizes > 1.0 are considered to represent about a 55% non-overlap, or dissimilarity of taxa, > 2.0 represents about 80% non-overlap or dissimilarity of taxonomic communities, > 3 represents more than 90% non-overlap or dissimilarity [49].

Additionally, a Canonical Analysis of the Principal Coordinates (CAP) was performed, using the PERMANOVA+ guidelines [48] [49], to provide a rigorous assessment of the distinctiveness of the microbial community compositions in the soils before and after the hurricane. Strong differences in the microbiome compositions between the different soil comparisons are indicated by CAP axis squared canonical correlations ≥ 0.7, and moderate differences are indicated by squared canonical correlations ≥ 0.5 to 0.69 [49]. To address question 3, we determined the differences in MPS of genera that were considered most important contributors to the soil communities, Mann-Whitney tests were conducted comparing the significance of the differences in MPS of genera with a MPS values ≥ 1.0% in one or more of the habitats studied, either in Pre or Post-Hurricane soil samples. We used a cutoff of p values < 0.05 to suggest significant differences between MPS values. These genera were then placed into one of the previously identified functional groups associated with the C and N cycle, if appropriate.

To address question 4, a distance-based linear model (DistLM) permutation test was implemented using PERMANOVA+ to determine if there were soil C and/or N-cycle variables that were significant predictors of the multivariate patterns of the soil bacterial and/or fungal community compositions associated with the different soil samples. We used the fourth root transformed resemblance Bray-Curtis matrices of the taxonomic data as response variables, the log (x + 1) transformed C and N data converted into Euclidean matrices as the predictor variables, and a step-wise selection process along with an AICc (Akaike’s Information Criterion Corrected) selection criterion and 9,999 permutations [48]. The AICc criterion is applied to handle situations where the number of samples (N) is small relative to the number (v) of predictor variables, where N/v < 40, as per Anderson et al.[48], and as was the case in this study.

## Results

### Differences in Soil C and N Data

The Lowland forest soils had greater levels of TN (*p* =0.043) in the Pre-Hurricane soil samples, and greater levels of NO_3_^-^ (*p* = 0.058), NH_4_^+^ (*p* = 0.012), Biomass C (*p* = 0.054), and Biomass C/TOC (*p* = 0.051) in the Post-Hurricane samples (Table 1). The Upland forest soils had greater levels of TOC (*p* = 0.022) and TN (*p* = 0.052) in the Pre-Hurricane soil samples, and greater levels of Biomass C (*p* = 0.033), Biomass C/TOC (*p* = 0.003), NO_3_^-^ (*p* = 0.010), and NH_4_^+^ (*p* = 0.012) in the Post-Hurricane soil samples (Table 1). The Riparian forest soils had greater levels of TOC (*p* = 0.018) and TN (*p* < 0.0001) in the Pre-Hurricane soil samples, and greater levels of Biomass C (*p* = 0.001), NO_3_^-^ (*p* = 0.038), and NH_4_^+^ (*p* = 0.012) in the Post-Hurricane soil samples (Table 1).

**Table 1.** The mean values of the soil environmental metrics from the 3 different forest types (Lowland, Upland, and Riparian), before and after Hurricane Otto, within the same forested area of the Maquenque National Wildlife Refuge in the Northern Zone of Costa Rica. The metrics analyzed were the Total Organic Carbon (TOC), Biomass C, Biomass C/TOC (Microbial Quotient), Total Nitrogen (TN), NH_4_^+^, and NO_3_^-^. The patterns of the significant differences in mean values between habitats, and ANOVA post-*hoc p* values are given.

| Habitat | TOC | Biomass C | Biomass C/TOC | TN | NH4 | NO3 |
| --- | --- | --- | --- | --- | --- | --- |
| Lowland Pre-Hurricane | 598.8 ± 23.7 | 410.8 ± 66.6 | 0.5 ± 0.09 | 48.2 ± 1.1 | 1.39 ± 0.21 | 8.6 ± 1.5 |
| Lowland Post-Hurricane | 594.2 ± 39.6 | 522.2 ± 69.0 | 0.88 ± 0.11 | 41.5 ± 1.2 | 5.1 ± 1.59 | 11.0 ± 2.0 |
| Upland Pre-Hurricane | 668 ± 28.4 | 351.4 ± 48.3 | 0.52 ± 0.05 | 52.8 ± 1.0 | 0.77 ± 0.30 | 18.3 ± 1.6 |
| Upland Post-Hurricane | 534.8 ± 27.0 | 629.6 ± 68.6 | 1.09 ± 0.08 | 44.8 ± 1.6 | 2.94 ± 0.28 | 24.6 ± 3.4 |
| Riparian Pre-Hurricane | 562.0 ± 29.7 | 427.8 ± 39.3 | 1.05 ± 0.13 | 50.6 ± 1.6 | 0.72 ± 0.18 | 14.0 ± 2.1 |
| Riparian Post-Hurricane | 525.2 ± 17.9 | 583.2 ± 60.8 | 1.01 ± 0.09 | 37.0 ± 1.4 | 2.01 ± 0.44 | 17.8 ± 1.8 |

### Differences in the Soil Bacterial Communities

The NGS of the soil eDNA successfully identified 1,342,185 bacterial DNA sequences, of which 538,121 could be categorized into 579 different bacterial genera. There were significant differences in the composition of the Total soil Bacterial communities found before and after the hurricane (Table 2) across the three forest types (Pseudo-F = 136.3, *p* = 0.0001). Specifically, the PERMANOVA pairwise results showed the magnitude of the differences were extremely large between the bacterial community compositions of the Pre- and Post-hurricane soil samples from the Lowland Forest sites (Pseudo-F = 327.2, *p* = 0.007), the Upland Forest sites (Pseudo-F = 202.7, *p* = 0.009), and the Riparian Forest sites (Pseudo-F = 185.7, *p* = 0.007). The CAP analysis (Figure 1a) further supported these findings, showing extremely strong separation between the soil bacterial community compositions of all Pre- and Post-Hurricane soil samples (CAP 1 R^2^ = 0.998, *p* = 0.0001).

**Table 2.**
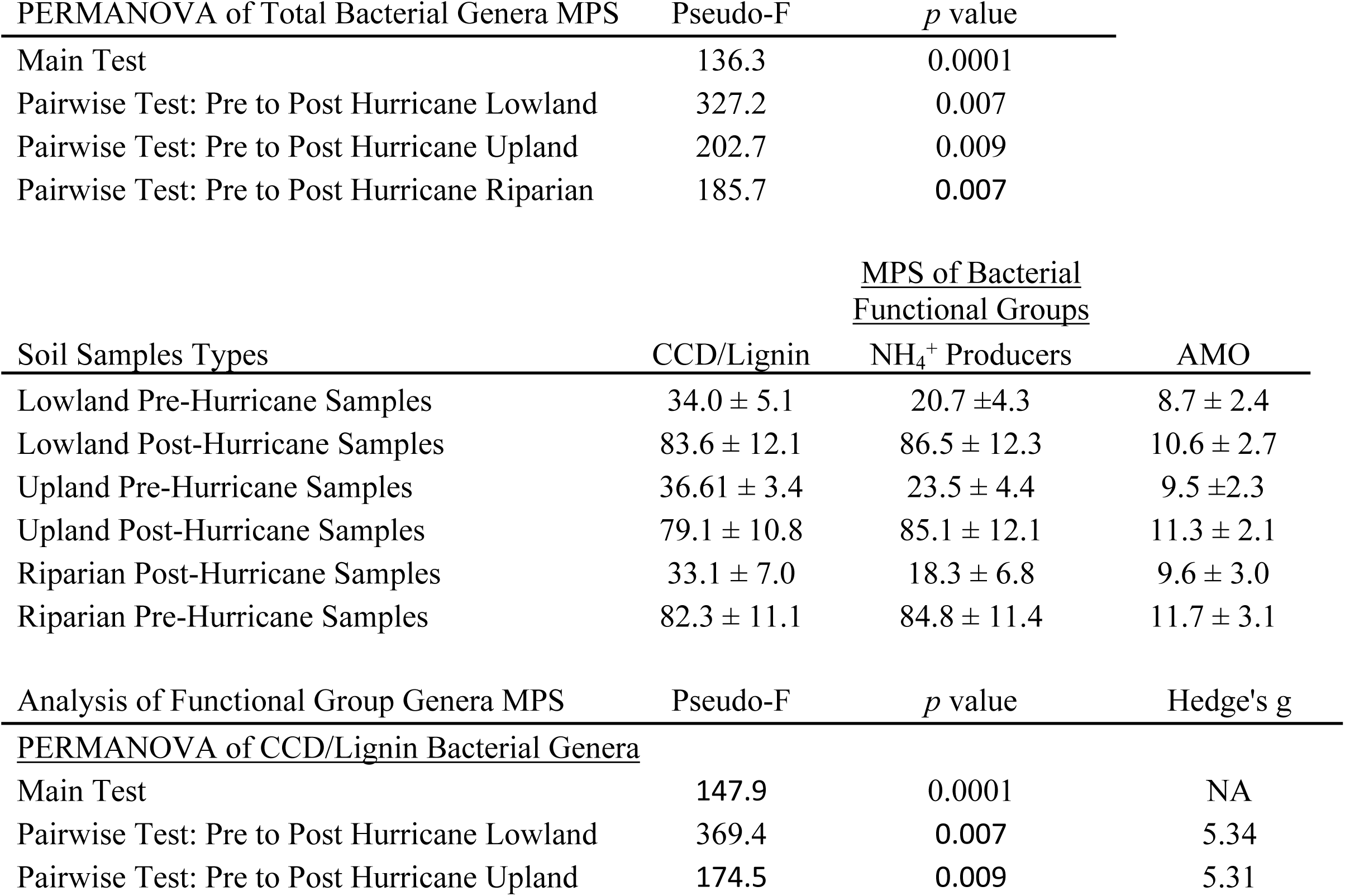
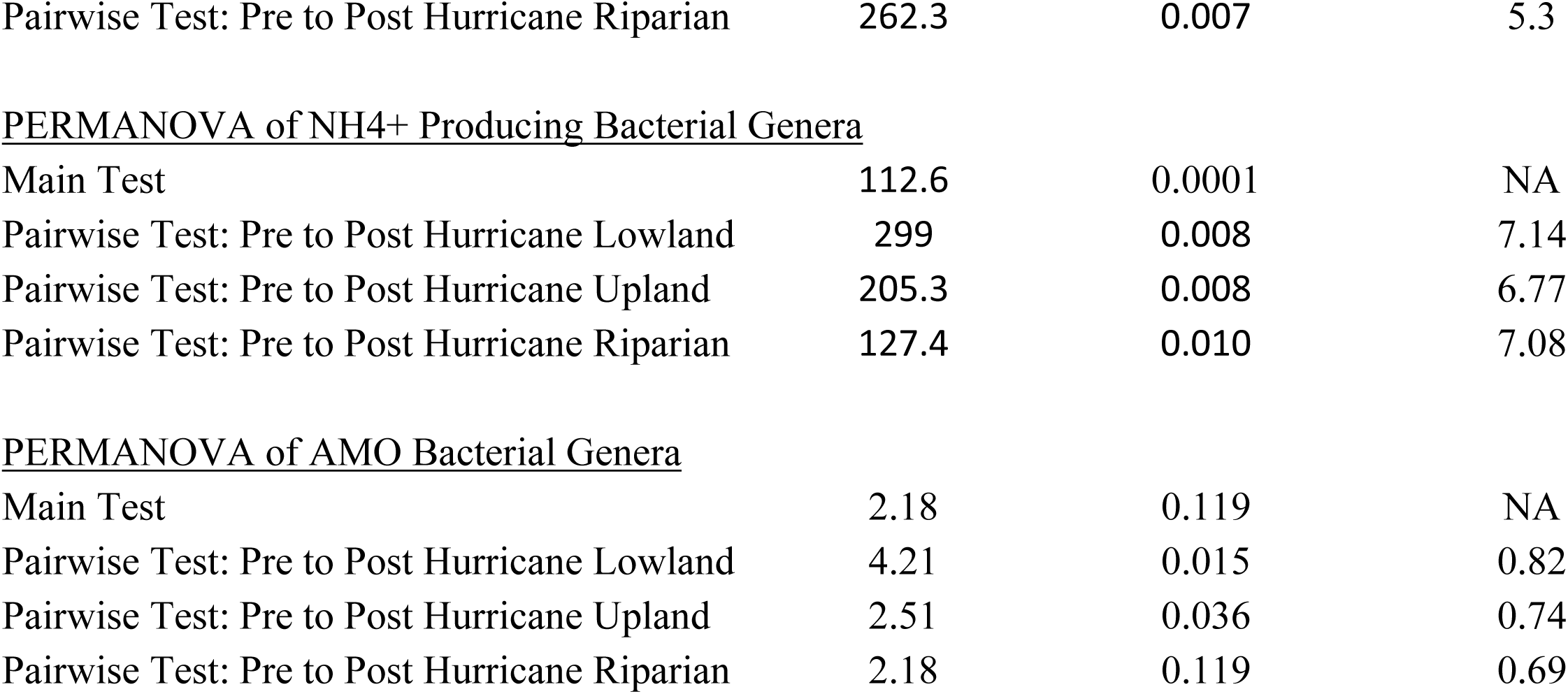
Results of the analysis of the mean proportion of DNA sequences (MPS) of different bacterial groups in forest soils before and after Hurricane Otto. The PERMANOVA Pseudo-F and *p* values are presented for the analysis of the differences in the MPS of the Total Bacterial Genera across the three forest types (Lowland, Upland and Riparian Forests) within the Maquenque National Wildlife Refuge in the Northern Zone of Costa Rica before and after Hurricane Otto. The MPS of the CCD/Lignin Degrading Bacterial Genera, the NH_4_^+^Producing Bacterial Genera, and the AMO Genera in the soils within three forest types (Lowland, Upland, and Riparian Forests) before and after the hurricane are also presented, along with the Main and Pairwise PERMANOVA Pseudo-F stat and *p* values, and Hedge’s *g* Effect size for each comparison.

| PERMANOVA of Total Bacterial Genera MPS | Pseudo-F | $p$ value | |
| --- | --- | --- | --- |
| Main Test | 136.3 | 0.0001 |  |
| Pairwise Test: Pre to Post Hurricane Lowland | 327.2 | 0.007 |  |
| Pairwise Test: Pre to Post Hurricane Upland | 202.7 | 0.009 |  |
| Pairwise Test: Pre to Post Hurricane Riparian | 185.7 | 0.007 |  |

| Soil Samples Types | <u>MPS of Bacterial Functional Groups</u> |  |  |
| --- | --- | --- | --- |
| | CCD/Lignin | $\text{NH}_4^+$ Producers | AMO |
| Lowland Pre-Hurricane Samples | 34.0 ± 5.1 | 20.7 ± 4.3 | 8.7 ± 2.4 |
| Lowland Post-Hurricane Samples | 83.6 ± 12.1 | 86.5 ± 12.3 | 10.6 ± 2.7 |
| Upland Pre-Hurricane Samples | 36.61 ± 3.4 | 23.5 ± 4.4 | 9.5 ± 2.3 |
| Upland Post-Hurricane Samples | 79.1 ± 10.8 | 85.1 ± 12.1 | 11.3 ± 2.1 |
| Riparian Post-Hurricane Samples | 33.1 ± 7.0 | 18.3 ± 6.8 | 9.6 ± 3.0 |
| Riparian Pre-Hurricane Samples | 82.3 ± 11.1 | 84.8 ± 11.4 | 11.7 ± 3.1 |

| Analysis of Functional Group Genera MPS | Pseudo-F | $p$ value | Hedge's $g$ |
| --- | --- | --- | --- |
| <u>PERMANOVA of CCD/Lignin Bacterial Genera</u> |  |  |  |
| Main Test | 147.9 | 0.0001 | NA |
| Pairwise Test: Pre to Post Hurricane Lowland | 369.4 | 0.007 | 5.34 |
| Pairwise Test: Pre to Post Hurricane Upland | 174.5 | 0.009 | 5.31 |
| Pairwise Test: Pre to Post Hurricane Riparian | 262.3 | 0.007 | 5.3 |
| <u>PERMANOVA of NH<sub>4</sub><sup>+</sup> Producing Bacterial Genera</u> |  |  |  |
| Main Test | 112.6 | 0.0001 | NA |
| Pairwise Test: Pre to Post Hurricane Lowland | 299 | 0.008 | 7.14 |
| Pairwise Test: Pre to Post Hurricane Upland | 205.3 | 0.008 | 6.77 |
| Pairwise Test: Pre to Post Hurricane Riparian | 127.4 | 0.010 | 7.08 |
| <u>PERMANOVA of AMO Bacterial Genera</u> |  |  |  |
| Main Test | 2.18 | 0.119 | NA |
| Pairwise Test: Pre to Post Hurricane Lowland | 4.21 | 0.015 | 0.82 |
| Pairwise Test: Pre to Post Hurricane Upland | 2.51 | 0.036 | 0.74 |
| Pairwise Test: Pre to Post Hurricane Riparian | 2.18 | 0.119 | 0.69 |

**Fig 1.**
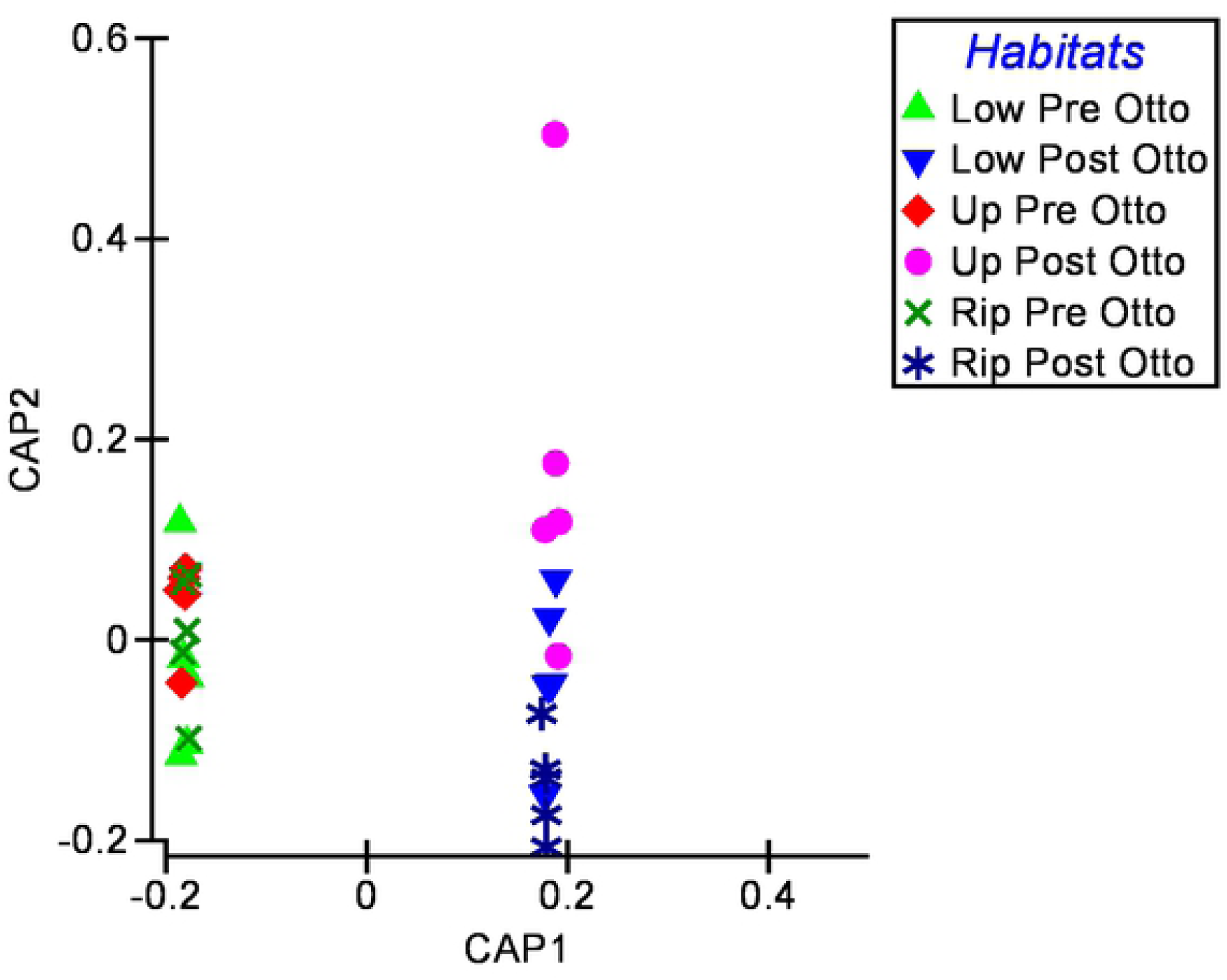
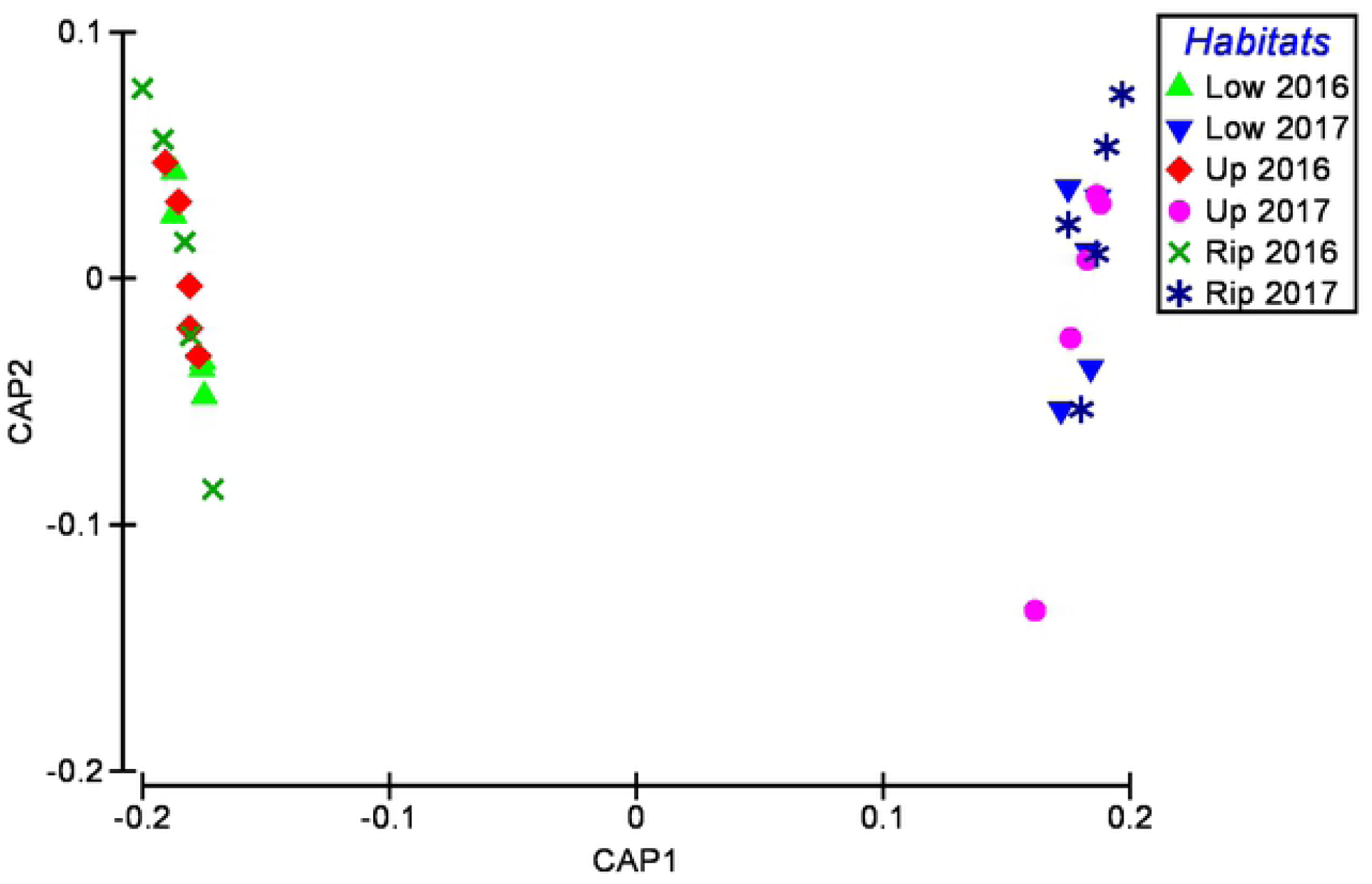
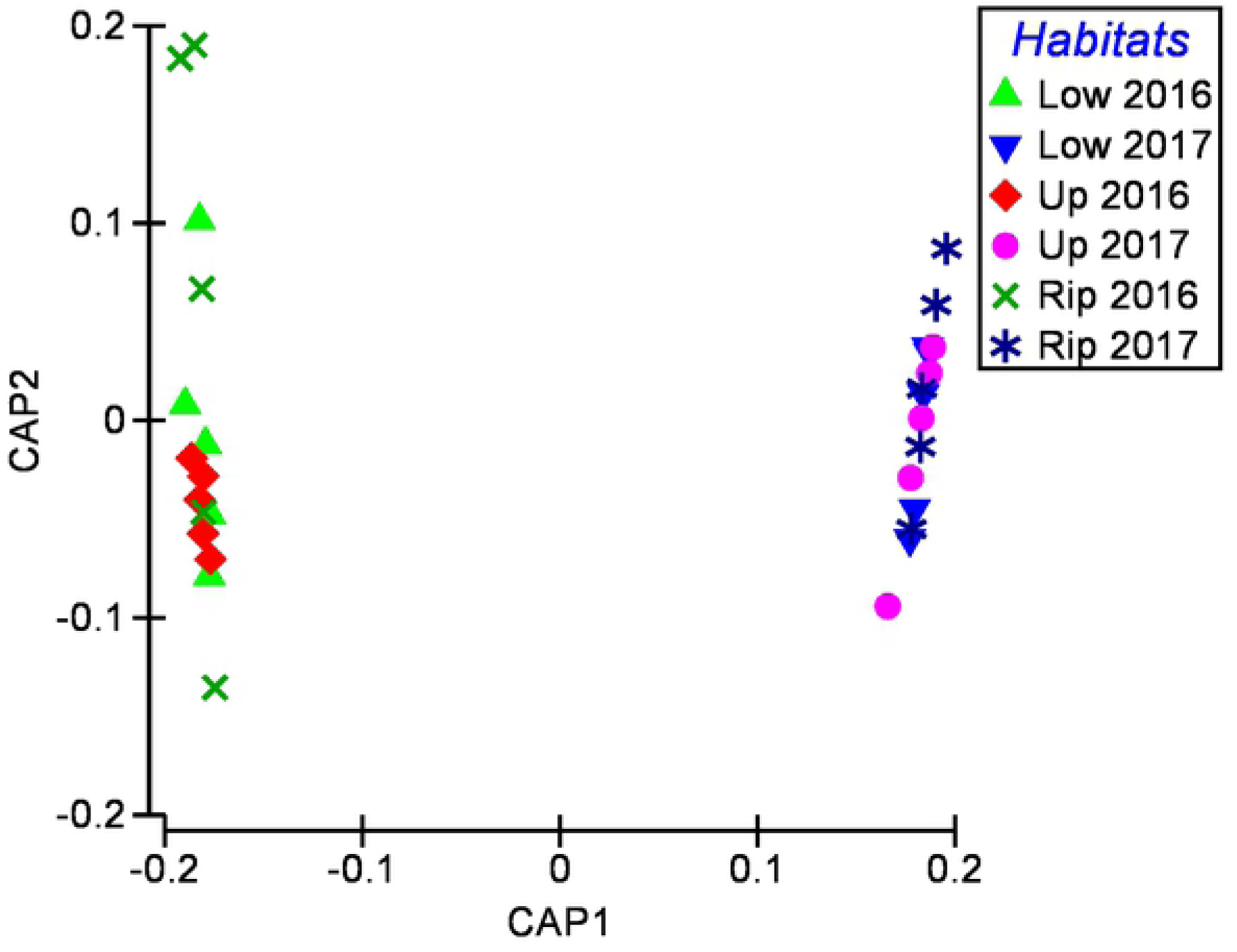
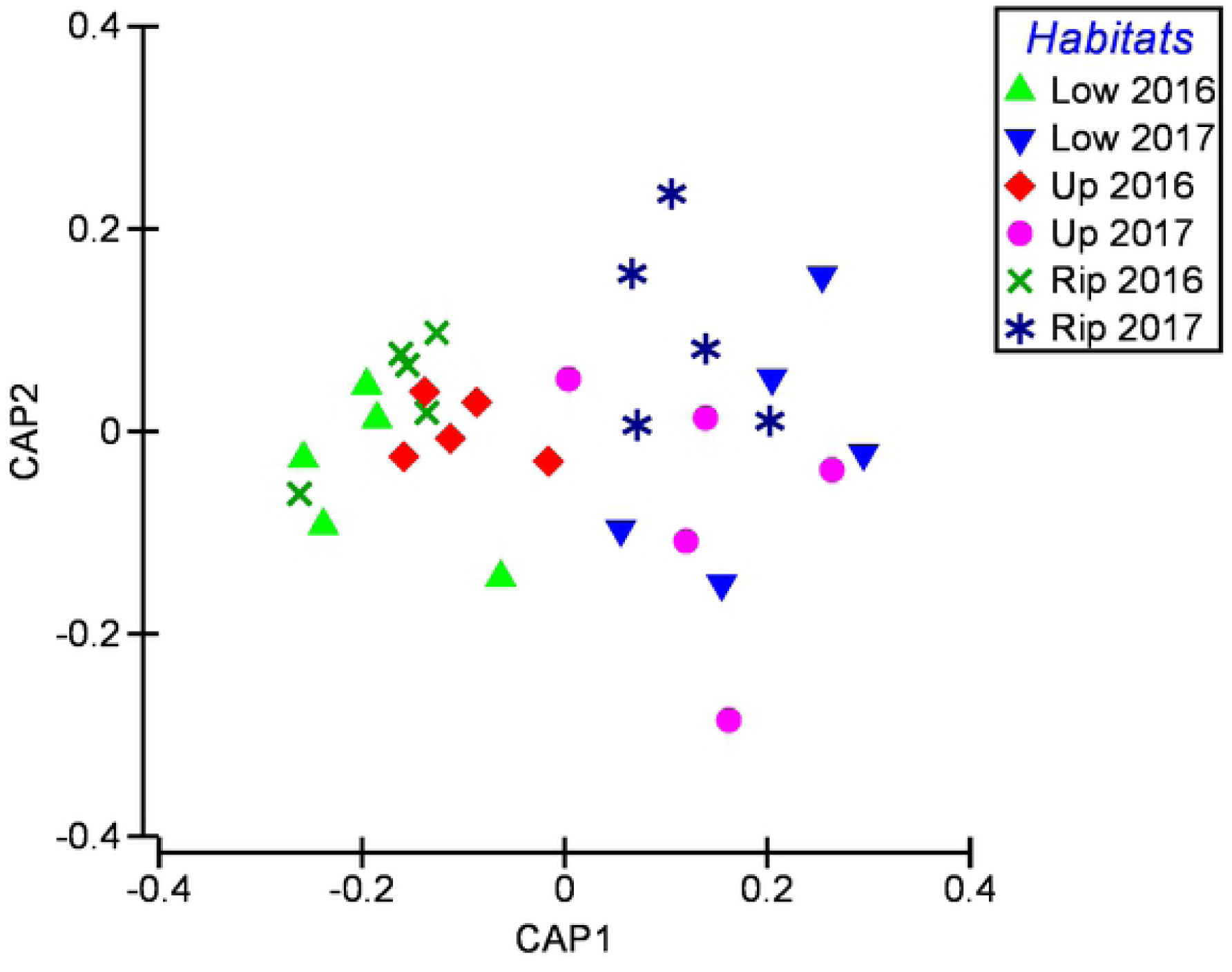
Results of the Canonical Analysis of Principal Coordinates (CAP) assessment of the differences in the patterns of the composition of the total bacterial genera (Fig 1a), the complex C and lignin degrading (CCD/Lignin) bacterial genera (Fig 1b), the NH_4_^+^ Producing bacterial genera (Fig 1c), and the AMO bacterial genera (Fig 1d) within soils from Lowland, Upland and Riparian Forest soils before and after Hurricane Otto hit these forests, which are within the Maquenque National Wildlife Refuge in the Northern Zone of Costa Rica.

There were also large differences in the MPS values of soil bacterial functional group communities before and after the hurricane (Table 2). There were far greater MPS values in the Post-Hurricane to Pre-Hurricane soil samples for the CCD/Lignin Degrading bacteria (Pseudo-F range = 174.5 to 369.4; *p* range = 0.007 to 0.009; Hedge’s *g* range = 5.30 to 5.34) and the NH_4_^+^ Producing bacteria (Pseudo-F range = 127.4 to 299.0; *p* range = 0.008 to 0.010; Hedge’s *g* range = 6.77 to 7.14). The Hedge’s *g* values shows there were 90-100% dissimilarity in composition of these two microbial communities in the soils before and after the hurricane in the two forest types. The MPS values of the AMO bacteria (Table 2) were also significantly greater in the Post-Hurricane to Pre-Hurricane soils from the Lowland and Upland forest types, although the magnitude of the differences were far less extreme than for the other two functional groups (Pseudo-F = 4.21 and 2.51; *p* = 0.015 and 0.036; Hedge’s *g* = 0.82 and 0.74, respectively). These Hedge’s *g* values suggests that there is about a 40-55% dissimilarity in the composition of this group in the soils of the two forest types. The CAP analyses (Figures 1b and 1c) also showed very strong separation in the structure of the CCD/Lignin Degrading and the NH_4_^+^ Producing bacterial communities between the Pre- and Post-Hurricane soil samples (CAP 1 R^2^ = 0.999, *p* = 0.0001; CAP 1 R^2^ = 0.998, *p* = 0.0001, respectively), but the model also reflected the lower level of separation in the structure of the AMO bacterial community in the soils between these three forest types (Figure 1d; CAP 1 R^2^ = 0.848, *p* = 0.0001).

The results of the Mann-Whitney tests (Table 3) conducted on the bacterial genera with MPS values > 1.0% within at least one habitat showed there were 11 genera (3 Acidobacteria Groups combined together) that were statistically different (*p* < 0.05) between the habitats. Extreme levels of difference in MPS values for these bacteria were evident by the Hedge’s *g* analysis, which showed far greater MPS levels of Acidobacteria Groups 1, 2, 3 (*g* = 10.87 to 58.51) and Verrumicrobia Spartobacteria (*g* = 4.15 to 6.07) in all Post-Hurricane soil samples, and that the Pre-Hurricane soil samples had greater MPS levels of *Isosphaera* (*g* = 4.32 to 8.41), *Koribacter* (*g* = 6.66 to 14.02), *Ktedonobacter* (*g* = 4.73 to 10.92), *Nitrospira* (*g* = 1.22 to 1.82), *Rhodoplanes* (*g* = 4.71 to 10.23), *Solibacter* (*g* = 8.42 to 28.94), and *Thermogemmatispora* (*g* = 3.35 to 10.53). Thus, all represent > large magnitude effects, and the differences in *Nitrospira* represent between 55% and 80% dissimilarity in the MPS of the genus within soils between before and after the hurricane, and the differences in all the others represent > 90% dissimilarity in the MPS of the genera.

**Table 3.** Results of the Mann – Whitney analyses showing the significant differences (*p* < 0.05) in the mean proportion of sequences per sample (MPS) of the genera that were present at > 1% MPS in the Lowland, Upland and Riparian Forest soils, before (Pre-Otto) and after (Post-Otto) Hurricane Otto hit the forests in November 2016. The mean MPS ± the standard deviation are given, as is the Hedge’s *g* Effect Size for the different comparisons.

| Differences in Bacterial<br>Genera (p < 0.05) |  | Lowland Forest Average MPS |  |  | Upland Forest Average MPS |  |  | Riparian Forest Average MPS |  |  |
| --- | --- | --- | --- | --- | --- | --- | --- | --- | --- | --- |
|  | Functions | Pre-Otto | Post-Otto | Hedge's *d* | Pre-Otto | Post-Otto | Hedge's *d* | Pre-Otto | Post-Otto | Hedge's *d* |
| Acidobacteria Groups 1, 2, 3 | CCD/Lignin, NH4+ Producers | 0.40 ± 0.12 | <u>71.12 ± 9.20</u> | 10.87 | 0.45±0.09 | <u>65.42 ± 2.64</u> | 34.78 | 0.31 ± 0.19 | <u>72.71 ± 1.74</u> | 58.51 |
| Verrucomicrobia Spartobacteria | CCD/Lignin, NH4+ Producers | 0.01 ± 0.01 | <u>3.53 ± 0.82</u> | 6.07 | 0.02 ± 0.01 | <u>5.35 ± 1.79</u> | 4.21 | 0.02 ± 0.02 | <u>3.16 ± 1.07</u> | 4.15 |
| Isosphaera | CCD/Lignin | <u>5.95 ± 0.96</u> | 0.24 ± 0.03 | 8.41 | <u>1.41 ± 0.31</u> | 0.25 ± 0.22 | 4.32 | <u>3.01 ± 0.72</u> | 0.35 ± 0.24 | 4.96 |
| Koribacter | CCD/Lignin | <u>20.52 ± 2.95</u> | 0.79 ± 0.16 | 9.44 | <u>23.39 ± 2.27</u> | 0.71 ± 0.29 | 14.02 | <u>24.33 ± 4.86</u> | 1.30 ± 0.54 | 6.66 |
| Ktedonobacter | unknown | <u>8.12 ± 1.02</u> | 0.22 ± 0.08 | 10.92 | <u>5.89 ± 1.07</u> | 0.26 ± 0.07 | 7.43 | <u>5.73 ± 1.69</u> | 0.08 ± 0.06 | 4.73 |
| Nitrospira | AMO | <u>6.07 ± 1.97</u> | 3.28 ± 0.91 | 1.82 | <u>5.23 ± 1.10</u> | 3.54 ± 1.62 | 1.22 | <u>8.54 ± 2.64</u> | 5.06 ± 2.44 | 1.37 |
| Rhodoplanes | NH4+ Producers | <u>14.23 ± 2.12</u> | 0.12 ± 0.02 | 9.41 | <u>16.75 ± 2.30</u> | 0.10 ± 0.07 | 10.23 | <u>12.69 ± 3.81</u> | 0.01 ± 0.01 | 4.71 |
| Solibacter | CCD/Lignin, NH4+ Producers | <u>25.39 ± 2.25</u> | 2.34 ± 0.74 | 8.42 | <u>28.77 ± 1.30</u> | 2.06 ± 0.12 | 28.94 | <u>25.78 ± 3.05</u> | 2.36 ± 0.39 | 8.65 |
| Thermogemmatispora | unknown | <u>4.33 ± 0.53</u> | 0.34 ± 0.08 | 10.53 | <u>4.44 ± 0.77</u> | 0.06 ± 0.05 | 8.03 | <u>5.10 ± 2.14</u> | 0.03 ± 0.04 | 3.35 |

| Differences in Fungal<br>Genera (p < 0.05) |  | Lowland Forest Average MPS |  |  | Upland Forest Average MPS |  |  | Riparian Forest Average MPS |  |  |
| --- | --- | --- | --- | --- | --- | --- | --- | --- | --- | --- |
|  | Functions | Pre-Otto | Post-Otto | Hedge's *d* | Pre-Otto | Post-Otto | Hedge's *d* | Pre-Otto | Post-Otto | Hedge's *d* |
| Calonectria | CCD/WRT/Lignin | <u>1.23 ± 0.59</u> | 0.37 ± 0.33 | 1.79 | <u>0.88 ± 0.43</u> | 0.51 ± 0.48 | 0.81 | <u>0.95 ± 0.29</u> | 0.53 ± 0.43 | 1.14 |
| Cryptococcus | CCD/WRT/Lignin | 12.88 ± 3.12 | <u>17.51 ± 8.56</u> | 0.81 | 1.26 ± 0.52 | <u>6.87 ± 3.75</u> | 2.1 | 10.89 ± 3.41 | <u>15.88 ± 5.69</u> | 1.06 |
| Geotrichum | CCD/WRT/Lignin | 2.73 ± 0.43 | <u>25.71 ± 6.74</u> | 4.82 | 1.02 ± 0.76 | <u>4.63 ± 1.50</u> | 3.04 | 14.29 ± 5.26 | <u>23.37 ± 10.47</u> | 1.11 |
| Gliocladiopsis | CCD/WRT/Lignin | 1.86 ± 1.09 | <u>4.64 ± 2.56</u> | 1.41 | 0.73 ± 0.38 | <u>3.31 ± 1.23</u> | 2.83 | 2.26 ± 1.36 | <u>10.62 ± 5.54</u> | 2.07 |
| Mortierella | CCD/WRT/Lignin | 4.78 ± 0.58 | <u>12.37 ± 6.87</u> | 1.56 | 8.88 ± 5.36 | <u>34.17 ± 16.32</u> | 2.08 | 1.24 ± 1.08 | <u>11.39 ± 8.31</u> | 1.71 |
| Myxocephala | unknown | <u>3.18 ± 1.97</u> | 1.22 ± 1.05 | 1.24 | <u>3.57 ± 1.55</u> | 1.17 ± 0.77 | 1.96 | <u>1.72 ± 0.56</u> | 1.05 ± 0.68 | 1.08 |
| Trichoderma | CCD/WRT/Lignin | <u>3.15 ± 1.81</u> | 0.78 ± 0.59 | 1.76 | <u>3.12 ± 0.96</u> | 0.53 ± 0.45 | 3.45 | <u>2.27 ± 1.80</u> | 0.27 ± 0.35 | 1.54 |
| Trichosporon | unknown | <u>41.3 ± 15.70</u> | 24.13 ± 11.86 | 1.23 | <u>46.48 ± 16.39</u> | 18.34 ± 7.00 | 2.23 | <u>35.08 ± 18.99</u> | 16.06 ± 6.67 | 1.33 |
| Xylaria | CCD/WRT/Lignin | <u>1.65 ± 1.11</u> | 0.72 ± 0.65 | 1.02 | <u>2.40 ± 1.42</u> | 0.28 ± 0.25 | 2.08 | <u>2.69 ± 1.17</u> | 0.57 ± 0.28 | 2.49 |

The significance and magnitude of the differences in the richness levels of bacterial communities (Table 4) were great across the soils of all three forest types before and after the hurricane. Specifically, the richness was greater in the Pre-Hurricane soils compared to the Post-Hurricane samples for the Total bacterial community (Pseudo-F range = 26.94 to 53.14; *p* range = 0.008 to 0.009; Hedge’s *g* range = 3.74 to 4.71), the CCD/Lignin degrading bacterial community (Pseudo-F range = 33.99 to 64.16; *p* range = 0.001 to 0.008; Hedge’s *g* range = 3.64 to 5.74), and the NH_4_^+^ Producing Bacterial community (Pseudo-F range = 55.95 to 118.59; *p* range = 0.006 to 0.008; Hedge’s *g* range = 3.93 to 5.59). The richness levels of the AMO bacterial community were significantly greater, although at a greatly reduced magnitude, in the Pre-Hurricane compared to Post-Hurricane soil samples for the Lowland forest and the Upland forest (Pseudo-F = 2.76 and 4.69; *p* = 0.053 and 0.009; Hedge’s *g* = 1.56 and 2.49, respectively).

**Table 4.**
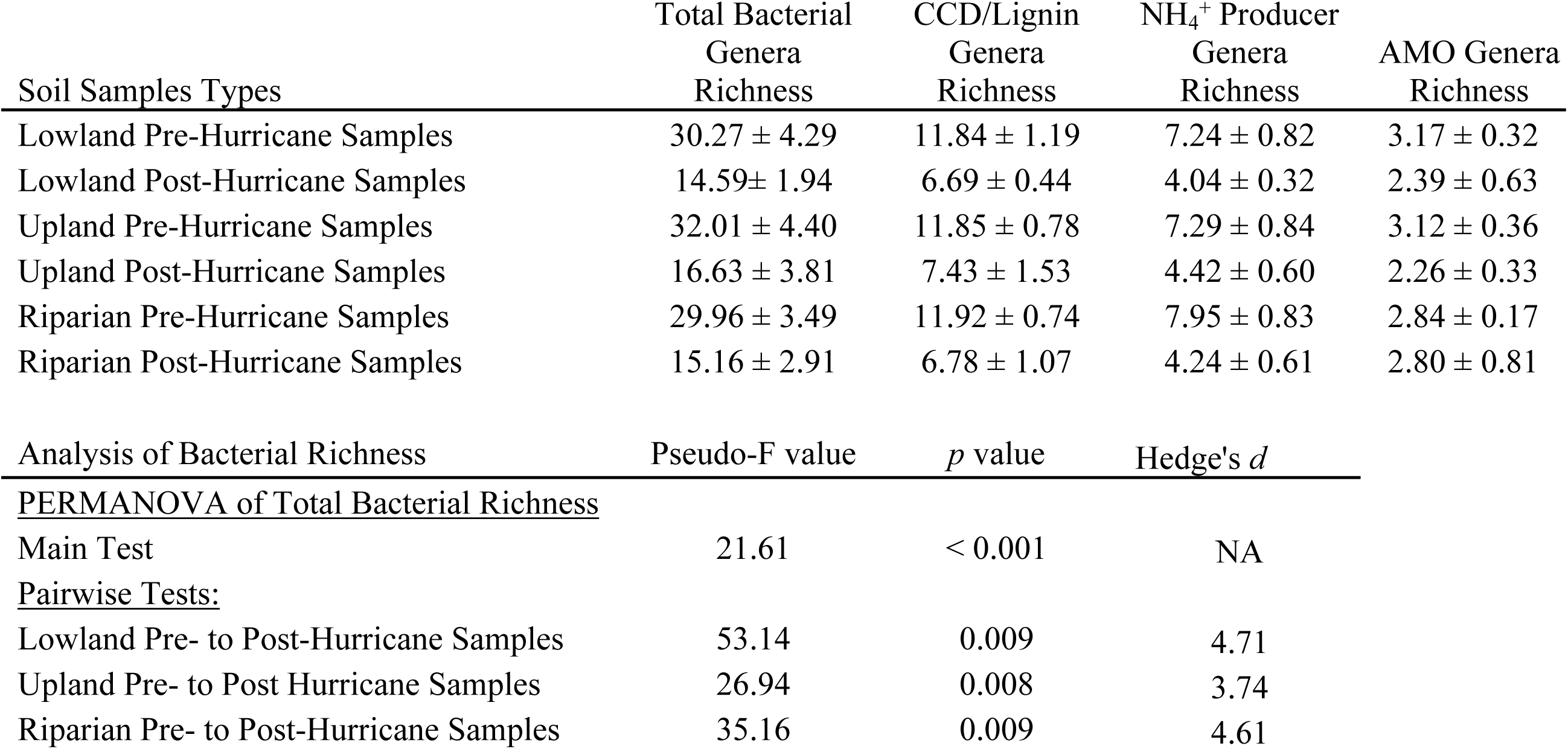
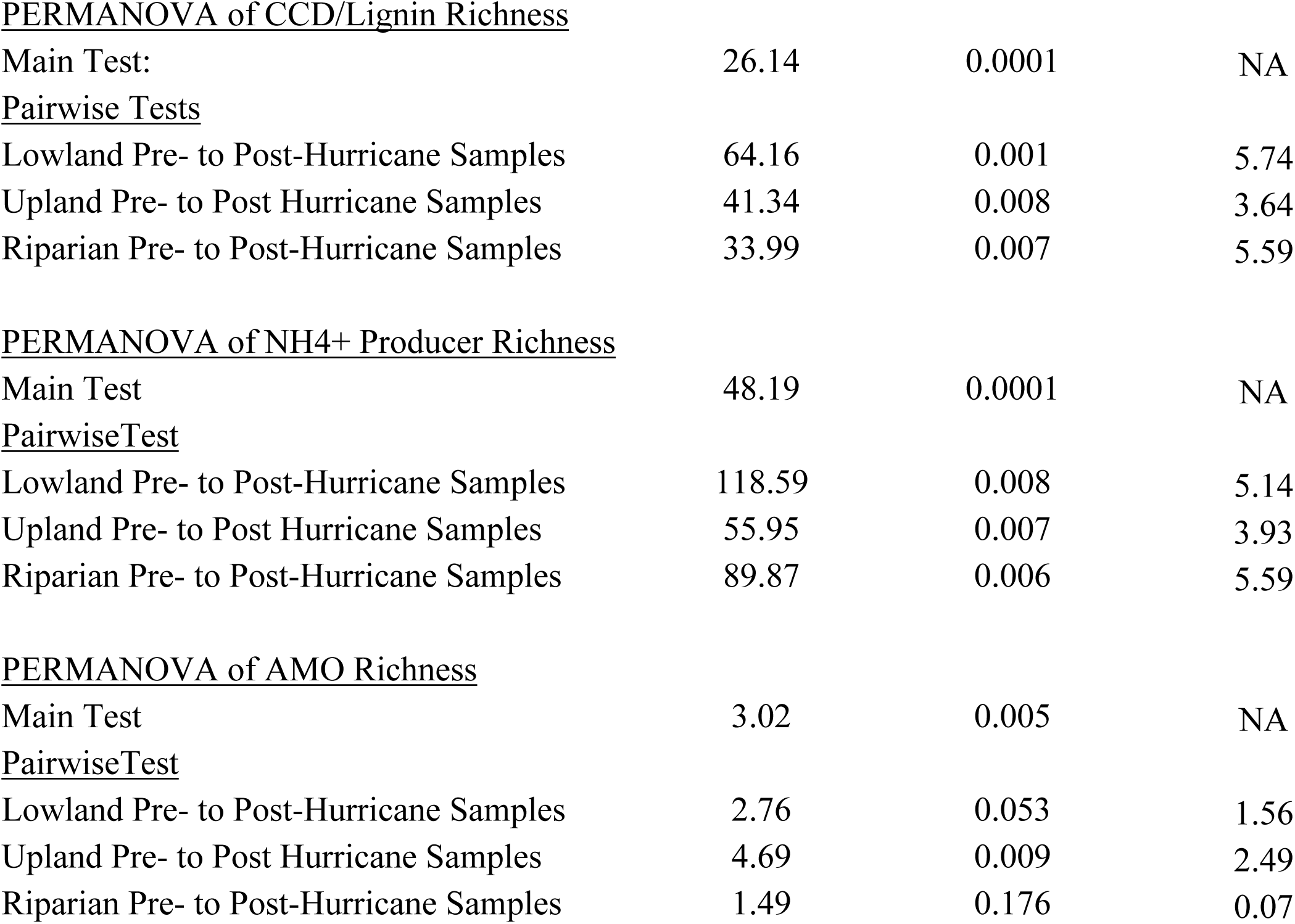
The mean richness (± standard deviation) of the Total Bacterial, CCD/Lignin degrading, NH_4_^+^ Producing, and AMO bacterial communities in the soils of the Lowland, Upland and Riparian Forest soils, before and after Hurricane Otto. In addition, the PERMANOVA Pseudo-F and *p* values, and the Hedge’s *g* Effect Size values for the analyses of the pairwise comparisons are given.

| Soil Samples Types | Total Bacterial<br>Genera<br>Richness | CCD/Lignin<br>Genera<br>Richness | $\text{NH}_4^+$ Producer<br>Genera<br>Richness | AMO Genera<br>Richness |
| --- | --- | --- | --- | --- |
| Lowland Pre-Hurricane Samples | 30.27 ± 4.29 | 11.84 ± 1.19 | 7.24 ± 0.82 | 3.17 ± 0.32 |
| Lowland Post-Hurricane Samples | 14.59 ± 1.94 | 6.69 ± 0.44 | 4.04 ± 0.32 | 2.39 ± 0.63 |
| Upland Pre-Hurricane Samples | 32.01 ± 4.40 | 11.85 ± 0.78 | 7.29 ± 0.84 | 3.12 ± 0.36 |
| Upland Post-Hurricane Samples | 16.63 ± 3.81 | 7.43 ± 1.53 | 4.42 ± 0.60 | 2.26 ± 0.33 |
| Riparian Pre-Hurricane Samples | 29.96 ± 3.49 | 11.92 ± 0.74 | 7.95 ± 0.83 | 2.84 ± 0.17 |
| Riparian Post-Hurricane Samples | 15.16 ± 2.91 | 6.78 ± 1.07 | 4.24 ± 0.61 | 2.80 ± 0.81 |
| Analysis of Bacterial Richness | Pseudo-F value | $p$ value | Hedge's $d$ | |
| <u>PERMANOVA of Total Bacterial Richness</u> |  |  |  |  |
| Main Test | 21.61 | < 0.001 | NA |  |
| <u>Pairwise Tests:</u> |  |  |  |  |
| Lowland Pre- to Post-Hurricane Samples | 53.14 | 0.009 | 4.71 |  |
| Upland Pre- to Post Hurricane Samples | 26.94 | 0.008 | 3.74 |  |
| Riparian Pre- to Post-Hurricane Samples | 35.16 | 0.009 | 4.61 |  |

PERMANOVA of CCD/Lignin Richness
|  |  |  |  |
| --- | --- | --- | --- |
| Main Test: | 26.14 | 0.0001 | NA |
| <u>Pairwise Tests</u> |  |  |  |
| Lowland Pre- to Post-Hurricane Samples | 64.16 | 0.001 | 5.74 |
| Upland Pre- to Post Hurricane Samples | 41.34 | 0.008 | 3.64 |
| Riparian Pre- to Post-Hurricane Samples | 33.99 | 0.007 | 5.59 |

PERMANOVA of NH<sub>4</sub><sup>+</sup> Producer Richness
|  |  |  |  |
| --- | --- | --- | --- |
| Main Test | 48.19 | 0.0001 | NA |
| <u>PairwiseTest</u> |  |  |  |
| Lowland Pre- to Post-Hurricane Samples | 118.59 | 0.008 | 5.14 |
| Upland Pre- to Post Hurricane Samples | 55.95 | 0.007 | 3.93 |
| Riparian Pre- to Post-Hurricane Samples | 89.87 | 0.006 | 5.59 |

PERMANOVA of AMO Richness
|  |  |  |  |
| --- | --- | --- | --- |
| Main Test | 3.02 | 0.005 | NA |
| <u>PairwiseTest</u> |  |  |  |
| Lowland Pre- to Post-Hurricane Samples | 2.76 | 0.053 | 1.56 |
| Upland Pre- to Post Hurricane Samples | 4.69 | 0.009 | 2.49 |
| Riparian Pre- to Post-Hurricane Samples | 1.49 | 0.176 | 0.07 |

The DistLM analyses showed that best predictor variable for variation in the MPS of the Total Bacterial community composition before the hurricane across the soils was the ToC levels (Table 5; AICc = 69.15, Pseudo-F = 3.54, *p* = 0.010), explaining 21.42% of the variation in the composition of the community. After the hurricane, the best predictor variables for the composition of the Total Bacterial community were Biomass C, NH_4_^+^, and NO_3_^-^ (Table 5; AICc = 71.60, Pseudo-F = 2.56, *p* = 0.045), explaining 36.67% of the variation in the composition of the community.

**Table 5.**
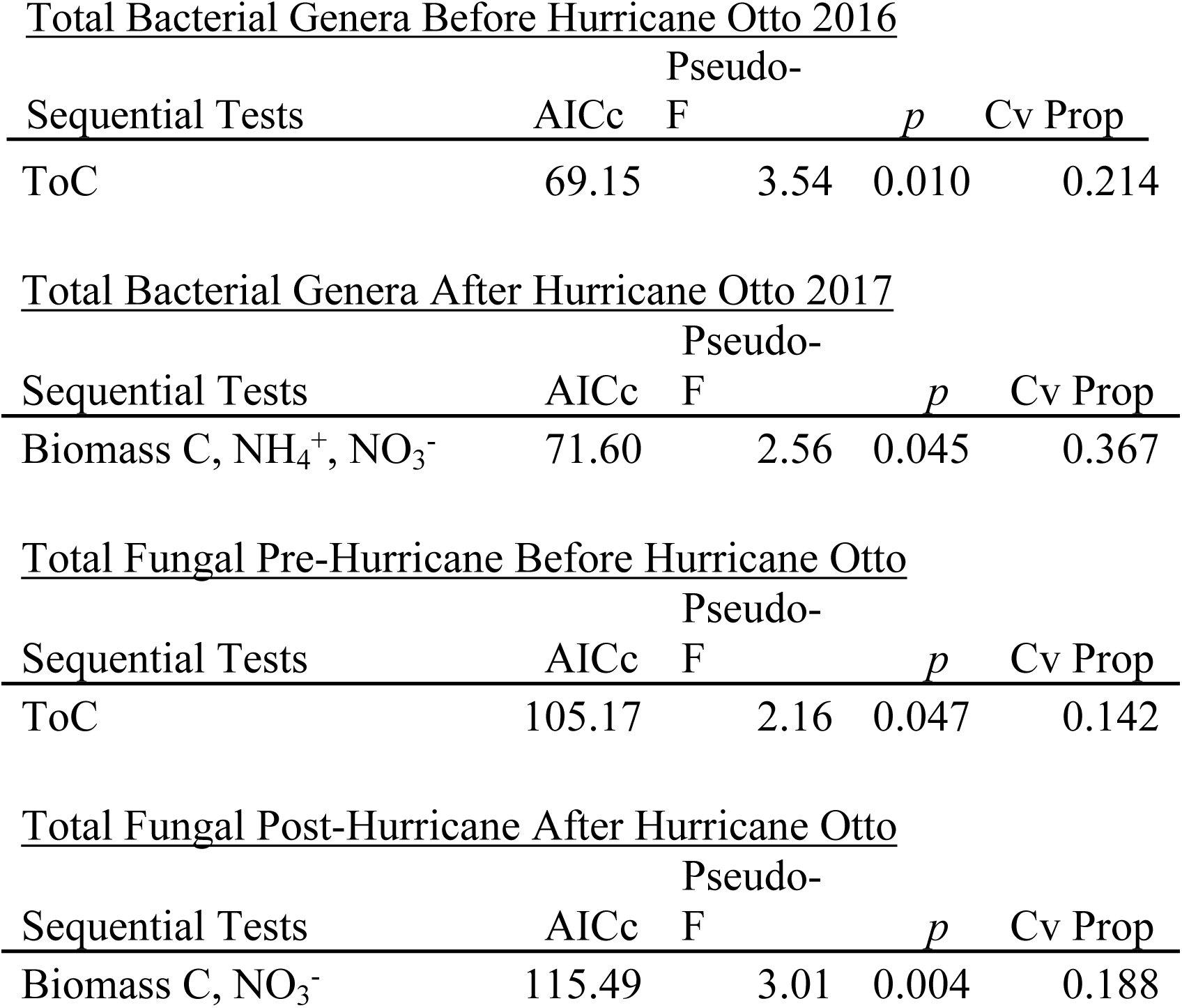
The distance-based linear modeling (DistLM) sequential tests describing the soil environmental variables that were the best predictors for the variation in the patterns in the composition of the Total Bacterial and the Total Fungal Genera communities, before and after Hurricane Otto, in the soils of three forest habitats within a primary forest area in the Maquenque National Wildlife Refuge in the Northern Zone of Costa Rica, using stepwise sequential tests following AICc criteria. The Pseudo-F and p values are given along with the cumulative proportion of the total variation in the microbial group composition (Cv Prop).

#### Differences in the Fungal Communities

The NGS of the soil eDNA successfully identified 1,114,085 fungal DNA sequences, of which 433,845 identified into 308different fungal genera. There were significant differences in the composition of the Total soil Fungal communities before and after the hurricane (Table 6) across the three forest types (Pseudo-F = 2.77, *p* = 0.0001). The PERMANOVA pairwise results showed the community compositions were different between the Pre- and Post-hurricane Lowland Forest soils (Pseudo-F = 2.16, *p* = 0.061), the Upland Forest soils (Pseudo-F = 4.22, *p* = 0.009), and the Riparian Forest sites (Pseudo-F = 2.30, *p* = 0.048). The CAP analysis (Figure 2a) confirmed these relationships, showing strong separation between the fungal community compositions of the Pre- and Post-Hurricane Upland Forest soil samples (CAP 1 R^2^ = 0.906, *p* = 0.0001), and between moderate and strong separation between the fungal community compositions in the Pre- and Post-Hurricane Lowland Forest and the Riparian Forest soil samples (CAP 1 R^2^ = 0.630, *p* = 0.0001). The DistLM analyses showed that best predictor variable for variation in the MPS of the Total Fungal community composition before the hurricane across the soils was the ToC levels (Table 5; AICc = 105.17, Pseudo-F = 2.16, *p* = 0.047), explaining 14.25% of the variation in the composition of the community. After the hurricane, the best predictor variable for the composition of the Total Fungal community was the level of Biomass C and NO_3_^-^ (Table 5; AICc = 115.49, Pseudo-F = 3.01, *p* = 0.004), explaining 18.78% of the variation in the composition of the community.

**Table 6.**
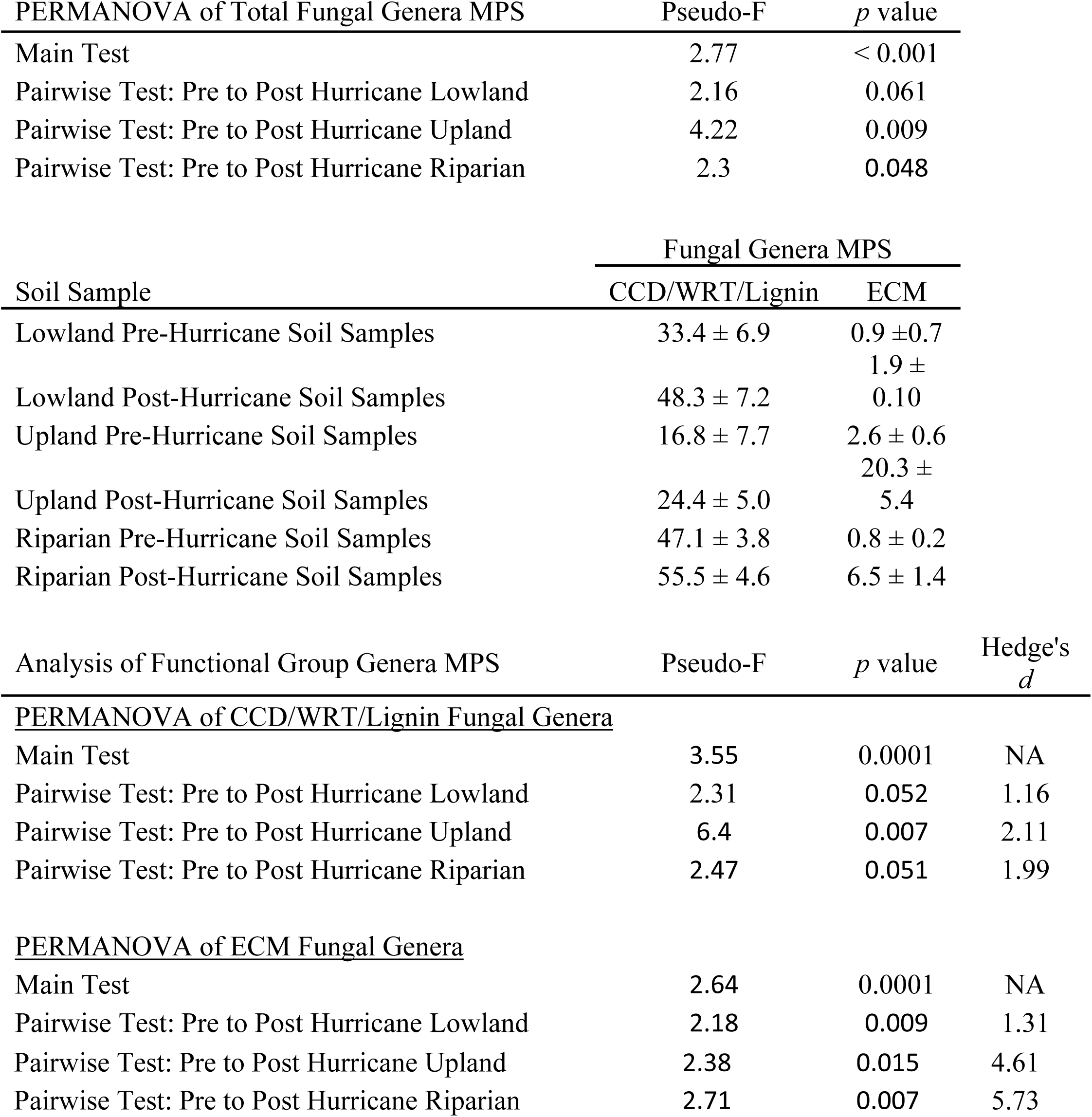
Results of the analysis of the mean proportion of DNA sequences (MPS) of different fungal groups in forest soils before and after Hurricane Otto. The PERMANOVA Pseudo-F and *p* values are presented for the analysis of the differences in the MPS of the Total Fungal Genera across the three forest types (Lowland, Upland and Riparian Forests) within the Maquenque National Wildlife Refuge in the Northern Zone of Costa Rica before and after Hurricane Otto. The MPS of the CCD/WRT/Lignin Degrading Fungal Genera and the ectomycorrhizal (ECM) fungal Genera in the soils within three forest types (Lowland, Upland, and Riparian Forests) before and after the hurricane are also presented, along with the Main and Pairwise PERMANOVA Pseudo-F stat and *p* values, and Hedge’s *g* Effect size for each comparison.

**Fig 2.**
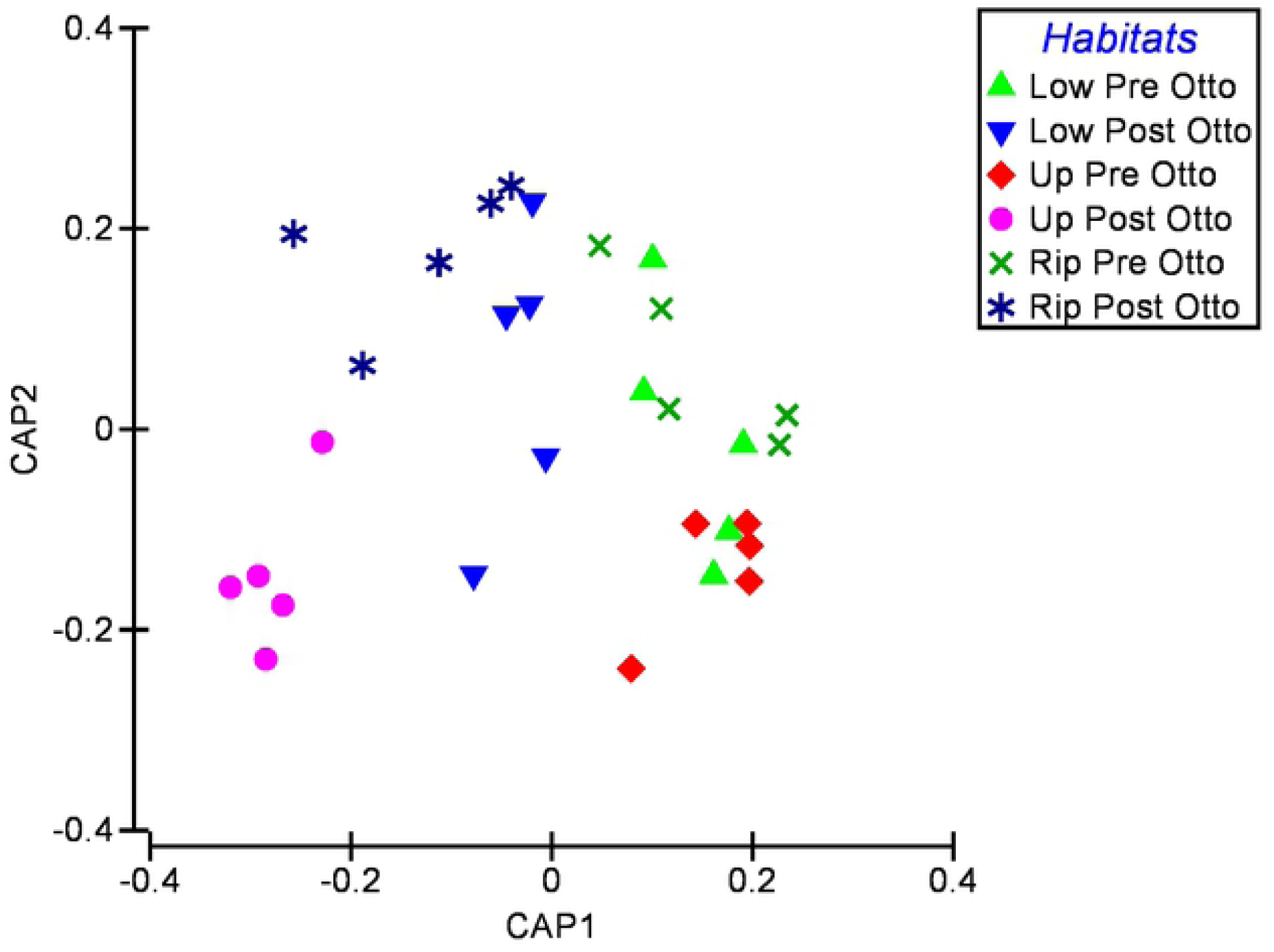
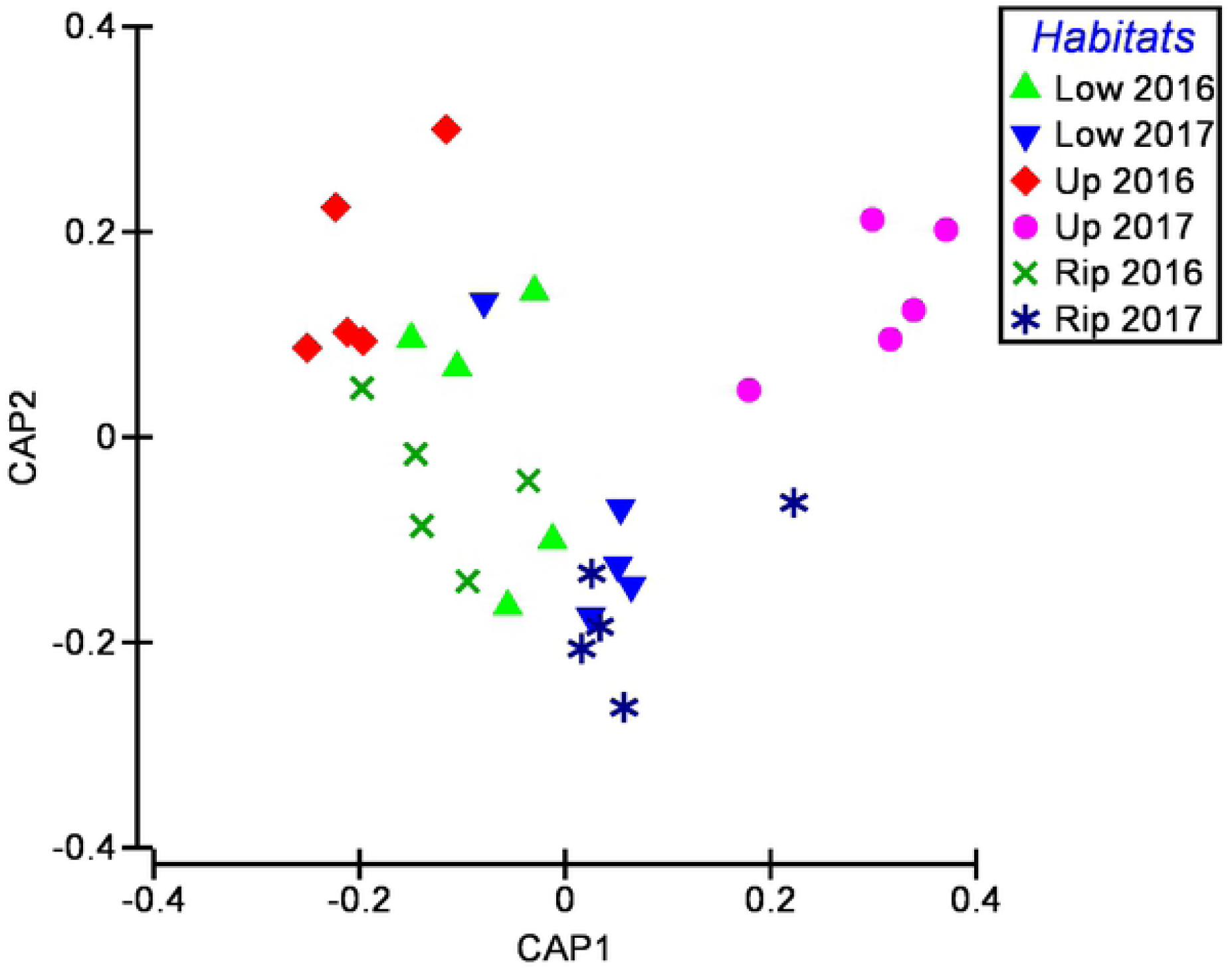
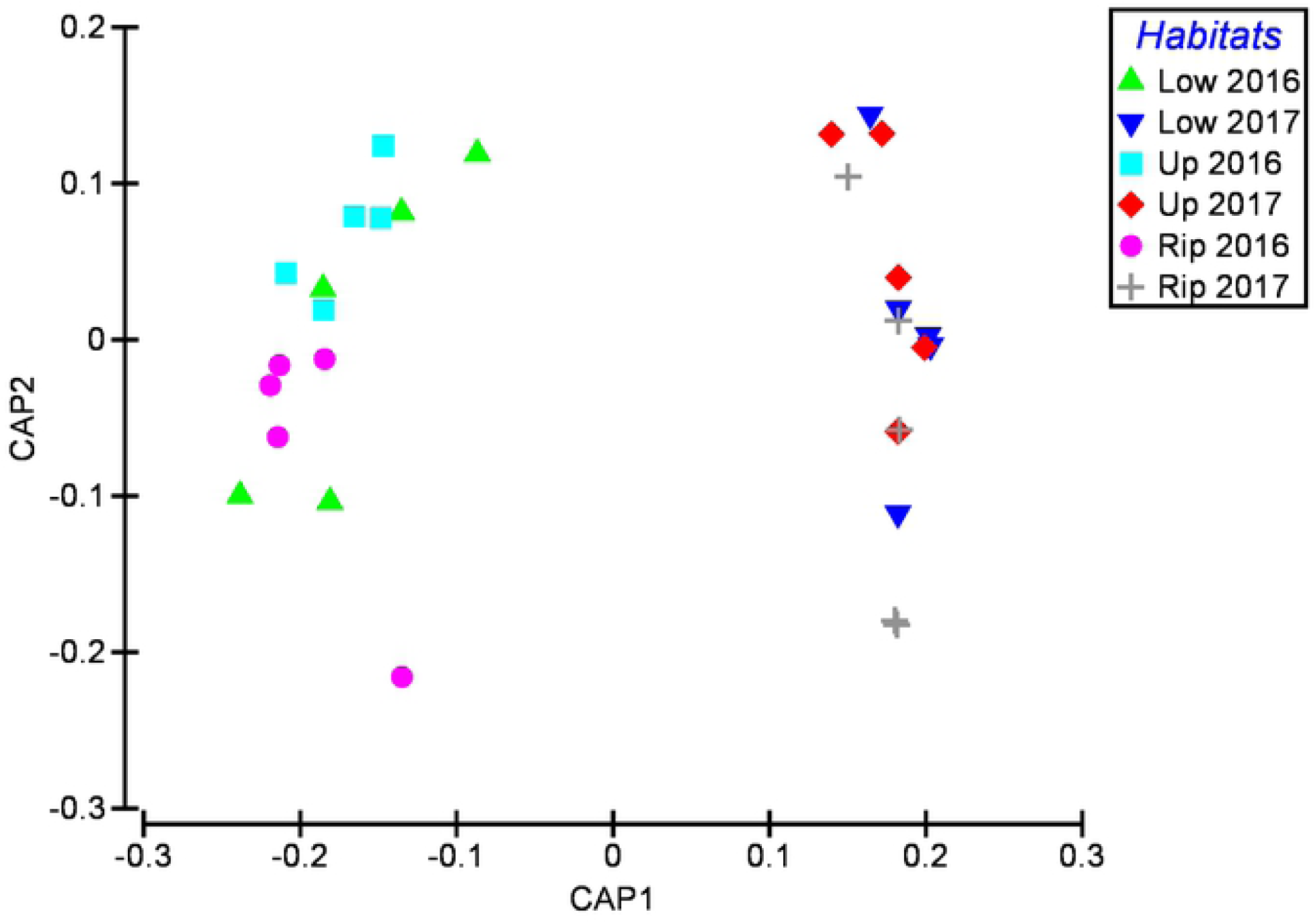
Results of the Canonical Analysis of Principal Coordinates (CAP) assessment of the differences in the patterns of the composition of the total fungal genera (Fig 1a), the complex C/wood rot/ and lignin degrading (CCD/WRT/Lignin) fungal genera (Fig 1b), the ectomycorrhizal (ECM) fungal genera (Fig 2c) within soils from Lowland, Upland and Riparian Forest soils before and after Hurricane Otto hit these forests, which are within the Maquenque National Wildlife Refuge in the Northern Zone of Costa Rica.

There were also large differences in the MPS values of the two soil fungal functional group communities before and after the hurricane (Table 6). The MPS values of the CCD/WRT/Lignin degrading fungal genera were greater in the Post-Hurricane to Pre-Hurricane soil samples from all three forest types (Pseudo-F range = 2.31 to 6.40; *p* range = 0.062 to 0.007; Hedge’s *g* range = 1.16 to 2.11), as were the MPS values of the community of ECM fungal (Pseudo-F range = 2.18 to 2.71; *p* range = 0.015 to 0.007; Hedge’s *g* range = 1.31 to 5.73). The CAP analyses (Figures 2b and 2c) were consistent with the PERMANOVA findings in that they showed strong separation in the community composition of the CCD/WRT/Lignin degrading fungal genera between the Pre- and Post-Hurricane soil samples from the Upland and the Riparian Forests (CAP 1 R^2^ = 0.884, *p* = 0.0001) and between moderate to strong separation of the composition of this fungal community in the Lowland Forest soil samples (CAP 1 R^2^ = 0.639, *p* = 0.0001). In addition, the CAP analyses (Figure 2c) showed very strong separation of the composition of the ECM genera community between the Pre- and Post-Hurricane soil samples for all three forest types (CAP 1 R^2^ = 0.975, *p* = 0.0118).

The results of the Mann-Whitney tests (Table 3) conducted on the fungal genera with MPS values > 1.0% within at least one habitat showed there were 9 genera with statistically different (*p* < 0.05) MPS values between the habitats. The MPS values of *Calonectria* (*g* = 0.81 to 1.79), *Myxocephala* (*g* = 1.98 to 1.96) *Trichoderma* (*g* = 1.54 to 3.45) *Trichosporon* (*g* = 1.23 to 2.23) and *Xylaria* (*g* = 1.02 to 2.49) were greater in the Pre-Hurricane soil samples of all three forest types, while the MPS values of *Cryptococcus* (*g* = 0.81 to 2.10), *Geotrichum* (*g* = 1.11 to 4.82), *Gliocladiopsis* (*g* = 1.41 to 2.83) and *Mortierella* (*g* = 1.56 to 2.08) were greater in the Post-Hurricane soil samples from all three forest types. Thus, all represent > large magnitude effects, and between 55% and > 90% dissimilarities in the MPS of the genera between the soils before and after the hurricane.

The patterns of the fungal richness were similar to that found for the bacterial richness in that the significance and magnitude of the differences in the richness levels of the fungal communities (Table 7). The fungal richness was greater in the Pre-Hurricane soils than the Post-Hurricane samples in all three forest types for the Total Fungal community (Pseudo-F range = 26.36 to 70.59; *p* range = 0.008 to 0.009; Hedge’s *g* range = 3.72 to 4.79), the CCD/WRT/Lignin degrading fungal community (Pseudo-F range = 17.53 to 56.57; *p* range = 0.008 to 0.009; Hedge’s *g* range = 4.01 to 4.09), and the ECM fungal community (Pseudo-F range = 101.10 to 225.36; *p* range = 0.007 to 0.009; Hedge’s *g* range = 7.37 to 9.45).

**Table 7.** The mean richness (± standard deviation) of the Total Fungal, CCD/WRT/Lignin degrading, and ECM fungal communities in the soils of the Lowland, Upland and Riparian Forest soils, before and after Hurricane Otto. In addition, the PERMANOVA Pseudo-F and *p* values, and the Hedge’s *g* Effect Size values for the analyses of the pairwise comparisons are given.

| Soil Samples Types | Total Fungal<br>Genera<br>Richness | WRT/CCD/Lignin<br>Genera Richness | ECM Genera<br>Richness |
| --- | --- | --- | --- |
| Lowland Pre-Hurricane Samples | 20.67 ± 4.22 | 8.39 ± 1.79 | 4.55 ± 0.71 |
| Lowland Post-Hurricane Samples | 4.65 ± 2.13 | 2.10 ± 1.24 | 0.14 ± 0.08 |
| Upland Pre-Hurricane Samples | 19.67 ± 2.72 | 9.19 ± 1.67 | 4.19 ± 0.72 |
| Upland Post-Hurricane Samples | 5.69 ± 4.56 | 3.75 ± 0.91 | 0.43 ± 0.04 |
| Riparian Pre-Hurricane Samples | 18.85 ± 4.56 | 6.51 ± 1.54 | 4.46 ± 0.50 |
| Riparian Post-Hurricane Samples | 4.82 ± 0.58 | 2.11 ± 0.20 | 0.92 ± 0.17 |

| Analysis of Fungal Richness | Pseudo-F value | *p* value | Hedge's *d* |
| --- | --- | --- | --- |
| <u>PERMANOVA of Total Fungal Richness</u> |  |  |  |
| Main Test | 22.05 | 0.0001 | NA |
| <u>Pairwise Tests:</u> |  |  |  |
| Lowland Pre- to Post-Hurricane Samples | 26.36 | 0.008 | 4.79 |
| Upland Pre- to Post Hurricane Samples | 70.59 | 0.009 | 3.72 |
| Riparian Pre- to Post-Hurricane Samples | 37.62 | 0.008 | 4.23 |
| <u>PERMANOVA of CCD/WRT/Lignin Fungal Richness</u> |  |  |  |
| Main Test: | 16.81 | 0.0001 | NA |
| <u>Pairwise Tests</u> |  |  |  |
| Lowland Pre- to Post-Hurricane Samples | 17.53 | 0.008 | 4.09 |
| Upland Pre- to Post Hurricane Samples | 29.38 | 0.008 | 4.05 |
| Riparian Pre- to Post-Hurricane Samples | 56.57 | 0.009 | 4.01 |
| <u>PERMANOVA of ECM Richness</u> |  |  |  |
| Main Test | 96.9 | 0.0001 | NA |
| <u>PairwiseTest</u> |  |  |  |
| Lowland Pre- to Post-Hurricane Samples | 101.1 | 0.008 | 8.73 |
| Upland Pre- to Post Hurricane Samples | 225.36 | 0.009 | 7.37 |
| Riparian Pre- to Post-Hurricane Samples | 217.87 | 0.007 | 9.45 |

## Discussion

Tropical forest hurricanes rapidly deposit large amounts of leaf litter and woody debris from the canopy to the forest floor [4], [5], [6], [7], which then results in short to long-term changes to the forest structure and forest ecological processes [4] associated with the rapid increase in the C and N nutrients [8], [9], [10], [11]. This increase in nutrients and enhancement of the ecological processes are thought to influence the soil microbial communities [12], [13], yet, very little is known about how hurricanes influence these communities. The current project is one of the few, to our knowledge, that used a “before and after” design to measure the effects that a naturally occurring hurricane had on the soil biotic and abiotic components within the first year, and within the same forest floor plots which had been studied the previous year. In this work, we provide evidence to confirm our predictions that there were increases in the soil inorganic N and Biomass C, and increases in the MPS, and changes in the community composition, of the general soil bacterial and fungal communities, and, more specifically, in the microbial communities associated with key C and N cycle processes in soil samples collected 3 months before and 8 months after Hurricane Otto hit the jungle.

There have been experimental studies that have shown the deposition of canopy material to the tropical forest floor, in a manner simulating the effects of a hurricane, result in nutrient pulses of more labile forms of C, N and phosphorus derived from the younger leaf litter from the canopy for about the first two years [4] [10] [14] [15] [16] [17]. This material is quickly processed within the first year [7] [11] [19], and is followed by utilization of the CWD deposited on the forest floor, resulting in a slower rate of processing activities [9] [11] [17] [18] [19] [20]. All these processes would be associated with the types of increases in inorganic N and Biomass C, and changes in the microbial communities that we observed in the forest soils after the hurricane [13] [36] [50] [51] [52] [53] [54].

In all three forest types for this study, the increases in the soil inorganic N and Biomass C after the hurricane were associated with overall changes in the bacterial and fungal community structure, as indicated by PERMANOVA and CAP analyses. The richness of the Total Bacterial and Total Fungal community were significantly reduced in the Post-Hurricane soils, suggesting that the excessive amounts of canopy leaf litter and CWD on the forest floor may be selecting for genera that become more dominant as they may be better able to rapidly process the newly deposited C and N-rich canopy material, resulting in enhanced processing of these materials. We provide several lines of evidence to support this theory. The DistLM analyses showed that the variables which suggest enhanced microbial activity were greater in the Post-Hurricane soils. Specifically, the best predictor variable for the variation in both the Total Bacterial and Total Fungal community compositions in the Pre-Hurricane soils was the level of ToC, while the best predictors for the composition of the Total Bacterial community in the Post-Hurricane soils were the levels of Biomass C, NH_4_^+^, and NO_3_^-^, and for the Total Fungal community they were Biomass C and NO_3_^-^. This could be the result of increases in the levels of microbial processing of nutrients occurring in the soils after the hurricane such that the values of the indicators of microbial activity (i.e., inorganic N and Biomass C levels) are greater after the hurricane hit. Furthermore, along with these observations, there were significant increases in the MPS values and decreases in richness of the CCD/Lignin degrading, NH_4_^+^ Producing and AMO bacterial genera, and the CCD/WRT/Lignin degrading and ECM fungal genera in the Post-Hurricane soil samples in all three forest types. As well, 4 of the 11 bacterial genera with MPS values > 1% that were found to have statistically greater MPS values in the Post-Hurricane samples (Acidobacteria Groups 1, 2, 3 genera, and Spartobacteria genera) actually composed about 70-75% of the bacterial MPS values in the Post-Hurricane samples, and were categorized as part of both the CCD/Lignin degrading and the NH_4_^+^ Producing bacterial functional groups. Furthermore, 4 of the 9 fungal genera with MPS values > 1% that were found to have statistically greater MPS values in the Post-Hurricane samples (*Cryptococcus, Geotrichum, Gliocladiopsis*, and *Mortierella*) composed about 50-60% of the fungal MPS values in the Post-Hurricane samples, and were categorized as CCD/WRT/Lignin degrading fungi. Collectively, these data support the idea that the canopy material that fell to the forest floor during the hurricane is stimulating specific changes to the soil microbial communities resulting in certain genera becoming more dominant that may be better adapted to processing the canopy material, and are associated with the observed increases in inorganic N and Biomass C.

Thus, in the current study, microbial functional groups and specific genera associated with enhanced utilization of the potentially enriched C and N nutrients, were present in greater amounts in Post-compared to Pre-Hurricane soil samples, and at reduced richness levels for all bacterial and fungal groups examined. These data suggest that these functional groups and specific genera within these groups are being selected for in the Post-Hurricane soils, and likely playing key roles in the processing and activation of the increased amounts of C and N nutrients in the soils, and subsequent incorporation of these materials into the food web. These results provide robust information and support for the findings of Lodge et al. [9] [14] and Cantrell et al [15] who demonstrated that differences occurred in the soil microbial community following experimental deposition of canopy material on the forest floor, without identifying taxonomic groups. We demonstrate support for their conclusions, and also provide the first information as to which bacterial and fungal genera are changing within Post-Hurricane in the soils, and the possible roles they are playing in the processing of the C and N nutrient cycles.

It would certainly be expected that changes in the quantity and quality of nutrient input from deposition of leaf litter and CWD due to a hurricane would cause alterations in soil microbial populations. For example, presumably, populations of the lignin degrading and complex C degrading bacteria and fungi, the wood rot fungi, the decomposer role of the ECM fungi (for example: [55] [56] [57] [58] [26] [27]), and many of the N cycle-associated bacteria (for example:[59] [25] [60] [61] [62] [63]) would all be enhanced following hurricane-deposited canopy material. It would also not be surprising that if these groups are being selected for after the hurricane, that the overall microbial richness might temporarily decrease, as these groups would be favored. All these changes in the soil microbial community would be expected to be associated with changes in decomposition activities, early shifts towards the breakdown of more labile forms of organic C, increased rates of CO_2_ release, changes in the rates and efficiency of use of soil organic of C and N, increased production of inorganic N, and general alterations to the biogeochemical and nutrient cycling processes within these soil ecosystems [36] [50] [51] [52] [53] [54]. The degree to which these activities occur within the belowground regions of the tropical forests post-hurricane will most certainly influence the rates of recovery of vegetation community diversity and health, and proper forest ecosystem functioning after the disturbance [12] [13].

## Conclusions

Hurricane Otto caused significant changes in the soil ecosystems of the three forests studied for this project, resulting in changes in the C and N cycle components and selecting for bacterial and fungal genera that may be better able to process the enhanced C and N materials. That said, the influences that hurricanes, in general, have on changes in the microbial community linked to the altered C and N processes post-deposition of litter and CWD still remains largely unknown, as are the longer-term patterns of soil ecosystem recovery following hurricane disturbances. Given the likelihood of an increasing frequency of hurricanes (or other extreme tropical weather events), and the apparent enhancement of C and N cycle processing by altered bacterial and fungal communities, determining how such major tropical forest disturbances influence the composition and functions of the forest soil ecosystems and ecological processes during recovery should be an area of significant interest, yet such studies are rare, receiving far less attention compared to that of the aboveground biota (Loreau, 2010). Future studies in these forests will begin to address this gap by long-term monitoring of the soils for changes in the microbial communities linked to changes in the C and N metrics over time to determine what soil ecosystem recovery looks like in these soils and how long it takes to get there.

## Acknowledgements

We would like to thank Vinzenz and Kurt Schmack, and all the staff members at the Laguna del Lagarto Lodge for their assistance and on-going support for our work. We would also like to thank the Pace University Dyson College Office of the Dean, the Dyson Faculty Research Grant Committee, and the Pace University Provost’s Office for their research support,. The authors declare they have no conflict of interest and that no animals or human subjects were used in this research.

